# A Robust Hierarchical Linear Model for Cryo-EM Map Analysis

**DOI:** 10.1101/2025.07.10.664269

**Authors:** I-Ping Tu, Shu-Cheng Zheng, Yu-Hsiang Lien, Szu-Han Lin, Po-Chun Lin, Wei-Hau Chang

## Abstract

Cryo-electron microscopy (cryo-EM) has become a central tool for determining the atomic structures of biological macromolecules, producing three-dimensional reconstruction maps that guide atomic model building. Current quantitative analyses of cryo-EM maps largely rely on simplified Gaussian signal models with fixed width parameters, limiting their ability to capture atom-specific signal characteristics directly from the maps. We propose a robust hierarchical linear (RHL) model for the statistical analysis of paired cryo-EM maps and atomic structures deposited in the EMDB and PDB. Within this framework, the logarithms of local voxel intensities surrounding each atom are modeled using a linearized Gaussian form, and atom-specific amplitude and width parameters are estimated through a hierarchical structure that pools information across atoms of the same type. The primary objective is to provide a statistically grounded framework for extracting quantitative atomic signal features from cryo-EM maps without imposing rigid physical assumptions on these parameters. To address contamination arising from overlapping atomic signals, spatially varying resolution, and experimental noise, we incorporate a data-adaptive weighting scheme based on minimum density power divergence estimation (MDPDE). This formulation preserves computational efficiency and ensures stable parameter estimation, while naturally facilitating the identification of deviated atomic profiles that may stem from model misspecification or local structural heterogeneity. Simulation studies demonstrate that the proposed method yields stable group-level parameter estimates even under substantial contamination. Applications to cryo-EM maps at multiple resolutions show that an atomic-resolution (1.25 Å) apoferritin map exhibits systematic atom-type differences in Gaussian amplitude and width, whereas a near-atomic-resolution map (2.20 Å) shows closer agreement with conventional uniform-width assumptions. Beyond structural biology, the proposed RHL framework provides a general statistical methodology for hierarchical data integration under heterogeneous noise, with potential applications in multi-center studies and federated learning.

## 1. Introduction

Over the past decade, single-particle cryo-electron microscopy (cryo-EM) has emerged as a central technique for determining the atomic structures of biological macromolecules. Cryo-EM reconstructs three-dimensional (3D) density maps from large collections of noisy two-dimensional (2D) projection images, and atomic models are subsequently built by fitting peptide chains into these maps. Each atomic model specifies the Cartesian coordinates of all atoms in a protein, and together with the corresponding cryo-EM map, is jointly deposited in the Protein Data Bank (PDB) and the Electron Microscopy Data Bank (EMDB).

Prior to 2013—the year often marked as the onset of the “resolution revolution”—the field of cryo-EM was dismissively termed “blob-ology” due to the limited resolution of reconstruction maps (Kühlbrandt, 2014; Saibil, 2017). During this era, the statistical toolkit for map analysis remained relatively sparse; because the data lacked atomic detail, there was little impetus to develop the “rich” quantitative frameworks required for sub-atomic signal quantification. Currently, even as resolution has reached the atomic scale, widely adopted metrics like the Q-score (Pintilie et al., 2020) and refinement pipelines such as PHENIX (Afonine et al., 2018) or REFMAC (Murshudov et al., 2011) continue to rely on legacy “uniform-width” Gaussian assumptions. Even AI-driven frameworks like ModelAngelo focus on coordinate optimization rather than the statistical quantification of signal parameters (Jamali et al., 2024). These standardized constraints, while ensuring stereochemical integrity, largely obscure the fine-grained, atom-specific signal variations that high-resolution data now potentially carry.

In this paper, we propose a Robust Hierarchical Linear (RHL) framework designed to extract and quantify atomic signal features directly from experimental cryo-EM maps. By utilizing a hierarchical structure to pool information across atoms of the same type, the model enables a robust statistical characterization of signal parameters. This framework shifts the focus from coordinate fitting and B-factor optimization (Rosenthal and Henderson, 2003) to direct signal quantification, offering the potential to identify chemically meaningful features that traditional pipelines are not designed to capture. While the Gaussian functional form remains the conventional kernel for modeling local intensity distributions (Afonine et al., 2018), these signal features have yet to be systematically quantified directly from reconstruction maps (Kirkland, 2020; Marques, Purdy and Yeager, 2019). It is therefore pertinent to investigate whether the Gaussian width parameter—intended to capture the spatial decay of the local signal—should vary across atom types, or if the uniform-width assumption is justified given the limitations of map quality.

Estimating a representative Gaussian parameter is nontrivial. At the atom level, signal overlap from nearby atoms (typically within 2 Å) elevates the local potential and breaks the radial symmetry expected from an isotropic Gaussian. At the group level, atoms located in flexible or solvent-exposed regions may deviate substantially due to increased mobility, compounded by spatially varying local resolution. These effects produce substantial deviations from the model and introduce contamination into the parameter estimates.

Fortunately, neighboring atomic contributions are not uniformly distributed around the target atom. Similarly, at the group level, flexible or heterogeneous regions generally constitute a limited fraction of the overall protein structure, which remains largely stable. Since these deviated intensities represent a limited fraction of the total observations, the hierarchical model can reduce the impact of contaminated voxels by incorporating a data-adaptive weighting scheme. This approach allows us to down-weight deviant data and recover a representative, group-level atomic signal.

We implement this deviation-downweighting scheme using weights derived from the Minimum Density Power Divergence Estimator (MDPDE) (Basu et al., 1998; Ghosh and Basu, 2013). The MDPDE minimizes the divergence between empirical and model distributions, yielding weighted estimators where individual weights are proportional to the likelihood raised to a positive power index. Because deviant observations yield smaller likelihood values, they are automatically assigned smaller weights. Therefore, larger power indices provide greater robustness by assigning smaller weights to deviant observations; as the power index approaches zero, the estimator converges to the Maximum Likelihood Estimator (MLE). To ensure map-specific calibration, we employ a cross-validation strategy to select these parameters, distinguishing between *α*_*r*_ for atom-level regression and *α*_*g*_ for estimating the group-level mean.

Furthermore, our methodological contribution extends beyond the application of weighted likelihoods by addressing both computational and structural limitations of existing robust methods. While earlier robust Bayesian approaches using density power divergence often lose the conjugate structure and incur heavy computational overhead (Ghosh and Basu, 2016), our framework incorporates MDPDE-derived weights directly into the hierarchical covariance structure, preserving the efficiency of the likelihood framework. Moreover, while existing MDPDE derivations typically restrict covariance to scalar or diagonal forms, we develop an explicit formulation for a fully unstructured, *p*-dimensional covariance matrix **Λ**. This generalization enables a more rigorous treatment of the complex dependence structures inherent in high-resolution cryo-EM data—such as the correlation between atomic amplitude and width—which simplified models are not designed to capture.

For illustration, we consider the backbone *C*_*α*_ atoms of alanine: each observed atom contributes a local neighborhood of map values to the group. In our model, a common mean parameter ***µ*** serves as the group-level representation, and each atom-specific parameter ***β***_*i*_ serves as an individual-level deviation:

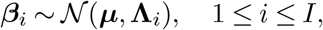

where *I* is the number of atoms in the group. This heteroscedastic formulation, by introducing **Λ**_*i*_, allows atom-level parameters to adapt to heterogeneous local map resolution and solvent exposure. For each atom, the neighborhood of local map values—after logarithmic transformation and denoted ***y***_*i*_—is modeled via linear regression of the Gaussian functional form. Because atoms of the same amino acid type are spatially distinct, we treat the ***β***_*i*_ as conditionally independent given the group-level parameters—an approximation that simplifies inference while retaining the dominant sources of variability in the data.

We first validate the method using simulated data and apply our approach to a suite of real-data maps spanning a range of resolutions from atomic (1.25 Å) to near-atomic levels (2.20 Å and 3.23 Å). While we provide a comparative analysis across this range, we focus our detailed findings on the atomic-resolution map of human apoferritin (PDB ID: 6Z6U, EMDB ID: 11103; resolution 1.25 Å). Notably, amide nitrogen consistently exhibits the lowest estimated amplitude and width across different amino acids. This observation suggests that the information content of atomic resolution cryo-EM maps may be higher than previously acknowledged, as the data appears to carry atom-specific information that the conventional uniform-width model fails to reflect. By moving beyond standardized constraints, our findings indicate that high-resolution cryo-EM maps can retain meaningful signal variation at the atomic scale, providing a foundation for future methodological and scientific investigations.

## 2. Method

### 2.1. Statistical Model and the Estimate

We define the cryo-EM map as a scalar field *y*(**r**) observed at discrete voxel locations within the 3D spatial domain ℝ^3^, where **r** = (*x, y, z*)^⊤^. This representation is theoretically grounded in the characterization of signal decay in reconstruction maps as a Gaussian fall-off in frequency space (Rosenthal and Henderson, 2003). Due to the Fourier duality of the Gaussian functional form, this frequency-domain decay corresponds to a localized Gaussian density in the spatial domain. Following established data extraction procedures (Pintilie et al., 2020; Afonine et al., 2018), we model the local signal around the *i*^*th*^ atom as a 3D Gaussian function:

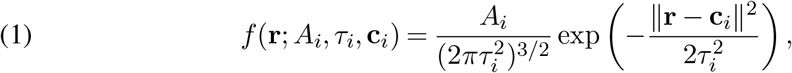

where **c**_*i*_ ∈ ℝ^3^ represents the Cartesian coordinates of the atom center, *A*_*i*_ is the amplitude, and *τ*_*i*_ is the width parameter (Afonine et al., 2018; Pintilie et al., 2020).

Figure 1 illustrates the data extraction process, where voxel intensities are sampled within a 1.5 Å radial distance of the atomic centers defined in PDB-6Z6U. Because the design matrix ***X*** depends solely on the radial distance *r*_*ij*_, the statistical model effectively operates as a one-dimensional regression. Importantly, the RHL analysis utilizes the full collection of 3D sampled voxel values directly, rather than the median-averaged radial profiles which are employed only for visualization purposes.

**FIG 1.**
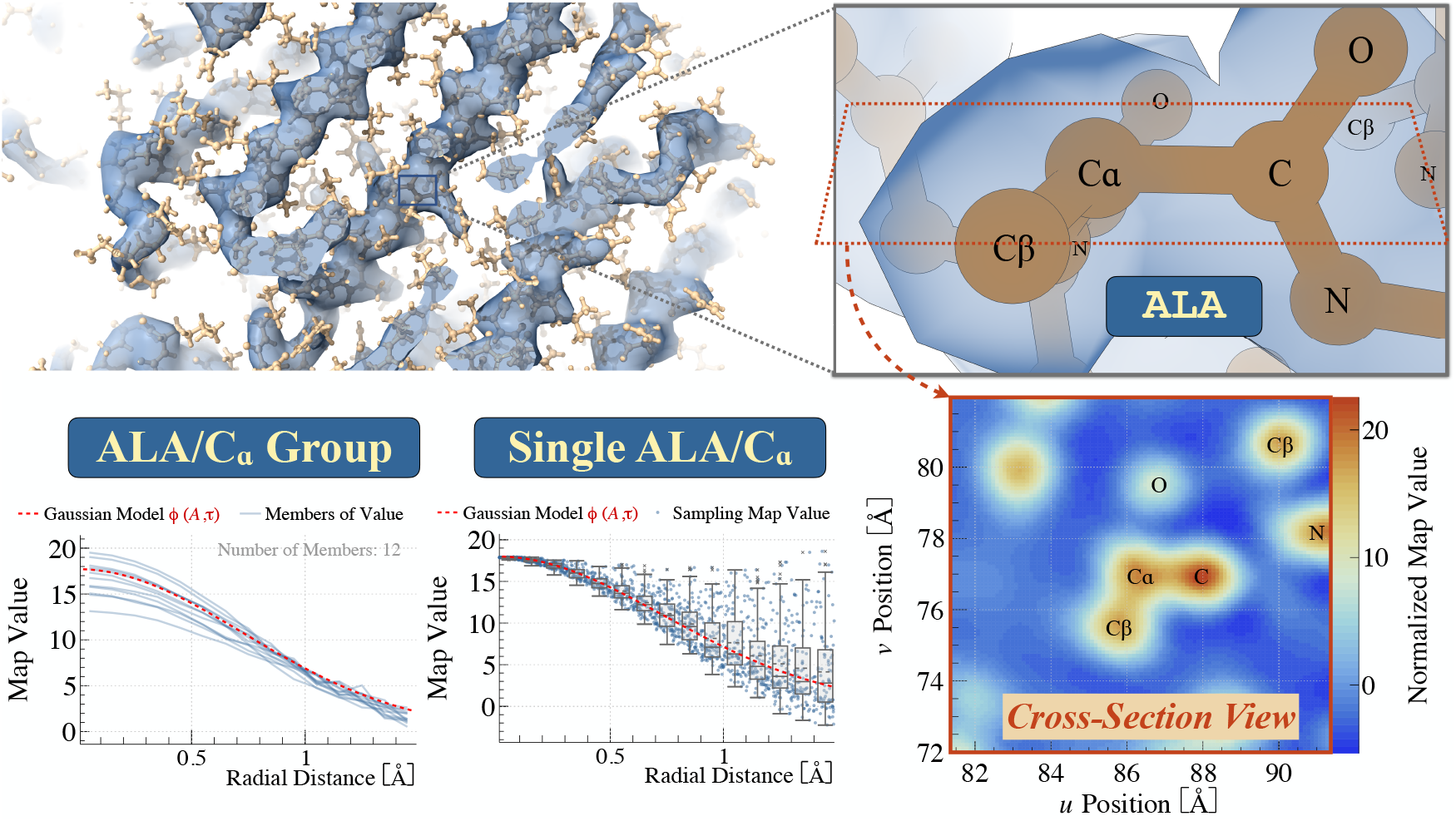
Illustration of Data Extraction from a Cryo-EM Map: We demonstrate the data extraction process using Cα atoms of alanine (ALA) residues in the cryo-EM map of human apoferritin (EMD-11103) and corresponding atomic coordinates (PDB ID: 6Z6U). Top-left: A Chimera representation where blue isosurfaces encode reconstructed map values and yellow circles indicate atomic model coordinates. Top-right: A magnified view of a single ALA residue highlighting its neighboring atoms. Bottom-right: The corresponding spatial intensity heatmap illustrating the cross-section of elevated intensities resulting from neighboring atoms, demonstrating that neighboring atoms contribute anisotropic perturbations to the local intensity field. Tricubic Interpolation: Using a tricubic interpolation method ((Afonine et al., 2018)), we sampled 1,500 map values within a 1.5 Å radial distance of each atomic center to extract the local atomic profile (see Section A.2 of the Supplementary Material (Tu et al., 2026) for the detailed sampling scheme). Bottom (“Single ALA/Cα”): The distribution of sampled intensities as a function of radial distance for an individual Cα atom. Bottom (“ALA/Cα Group”): Twelve curves representing independent ALA repeats within the apoferritin unit. Each curve represents a one-dimensional radial profile derived by computing the median map intensity at discrete radial intervals. The close adherence of these median-value curves to a Gaussian profile validates the selection of the Gaussian functional form as a representative model for atomic signals.

It is important to note that various formulations exist for the numerical representation of 3D Gaussian density functions, differing primarily by the normalization factor. Without the factor 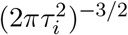 in Equation (1), *A*_*i*_ would refer to the peak intensity; however, by including this factor, *A*_*i*_ represents the **total intensity**. This choice ensures that *A*_*i*_ captures the integrated signal strength, which is more robust for quantifying atom-specific signal features.

To facilitate linear estimation, we transform the map values sampled at discrete grid points **r**_*ij*_. Let ∥*r*_*ij*_ = **r**_*ij*_ − **c**_*i*_∥ denote the radial distance from the *j*^*th*^ sampling point to the center of atom *i*. By applying a logarithmic transformation to the local voxel intensities, we obtain the following linear regression model:

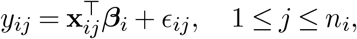

where *y*_*ij*_ represents the log-transformed intensity. The derivation is provided in Section A.1 of the Supplementary Material (Tu et al., 2026). As detailed in A.2 of the Supplementary Material (Tu et al., 2026), we sample 1,500 map values within the 1.5 Å radius to capture the primary signal while reducing interference from neighboring atoms. The design vector is defined as 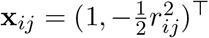, and the coefficient vector ***β***_*i*_ = (*β*_*i*0_, *β*_*i*1_)^⊤^ relates to the physical parameters as follows:

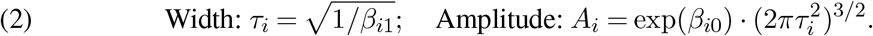

We extend this to a hierarchical framework by introducing an additional layer for the parameters ***β***_*i*_:

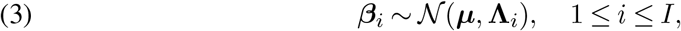

where ***µ*** denotes the group-level mean and **Λ**_*i*_ is an atom-specific covariance matrix. This heteroscedastic formulation allows the model to adapt to varied local environments, such as flexible loops or solvent-exposed regions. Stacking the observations for atom *i*, let 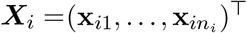 and 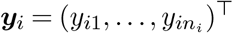. The regression model for each atom *i* can then be expressed in vector form as:

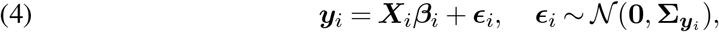

where ***y***_*i*_ is a vector of length *n*_*i*_. Notably, this heteroscedastic framework allows for unequal variances across observations, providing the necessary flexibility to accommodate the spatially varying noise and structural heterogeneity characteristic of cryo-EM reconstruction maps. Equations (3) and (4) collectively define a hierarchical, two-level statistical frame-work for analyzing atomic Gaussian parameters within specific amino acid categories. This heteroscedastic structure provides the necessary foundation for integrating robust estimation properties—specifically the downweighting of deviant intensities, as discussed in Section 1. Consequently, the MDPDE, which utilizes a data-adaptive weighting scheme, can reduce the influence of these deviations to recover a representative group-level signal. A detailed derivation of the weight construction using the MDPDE framework is provided in Section 2.2.

**TABLE 1.**
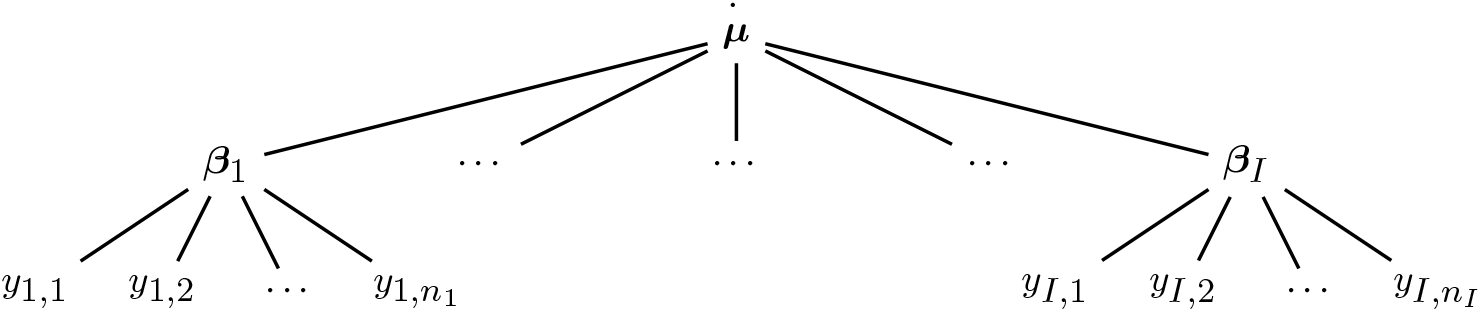
Model Structure: Estimate for Cα of ALA, I = 12.

We focus on the backbone atoms, alpha carbon (*C*_*α*_), carbonyl carbon (*C*), and nitrogen (*N*), as they provide the most reliable features for fitting atomic models to cryo-EM maps. To illustrate, we consider the alanine (ALA) residues in human apoferritin (PDB 6Z6U).

While the protein contains 288 ALA repeats, its 24-fold symmetry reduces these to *I* = 12 independent observations. For consistency, we assume *n*_*i*_ = *n* sampling points for each atom. Within this Robust Hierarchical Linear (RHL) framework, ***µ*** represents the group-level mean for a specific atom type (e.g., *C*_*α*_) within a specific amino acid category (e.g., ALA), while ***β***_*i*_ captures individual variations. Notably, we do not restrict the covariance matrix **Λ**_*i*_ to a diagonal form. This allows the model to faithfully represent the complex dependence structures between the amplitude and width parameters.

This two-level structure simplifies the three-level hierarchical model in Gelman et al. (2013), providing the heteroscedastic flexibility to down-weight deviant intensities through marginal likelihood estimation. We demonstrate that this framework can be applied to a portfolio of maps—ranging from 1.25 Å (6Z6U) and 2.20 Å (6DRV) to 3.23 Å (9GXM).

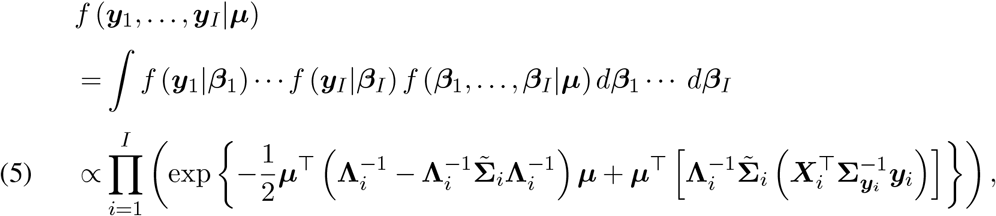

where 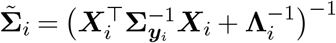.

Let *Q*(***µ***) be the exponent of Equation (5), we have

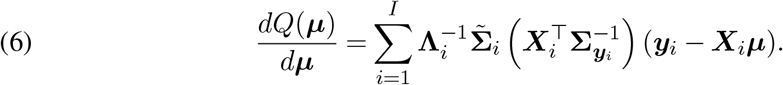

Then, the MLE by solving 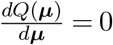 is as follows.

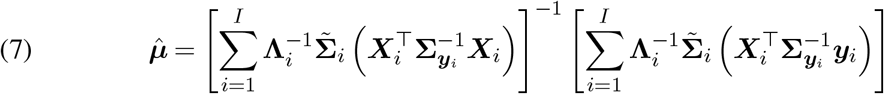

Here, in deriving 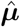, we assume that the covariance matrices 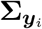 and **Λ**_*i*_, as specified in Equations (3) and (4), are known. In Section 2.2, we use the minimum density power divergence estimate (Basu et al., 1998; Ghosh and Basu, 2013) to construct the covariance matrices 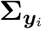 and **Λ**_*i*_. Detailed derivations are provided in Sections A.3–A.5 of the Supplementary Material (Tu(2026)).

### 2.2. Robust Estimates Formulated as Weighted Estimates

Basu et al. Basu et al. (1998) introduced the minimum density power divergence as a robust estimation framework where estimates can be expressed as weighted averages of the observed data. In this approach, each observation contributes to the estimate through a weight proportional to the density function *f*^*α*^, where *α >* 0 is a predetermined tuning parameter. This parameter *α* plays a crucial role in balancing robustness against data contamination with statistical efficiency. Weighting schemes have been demonstrated as an effective strategy for robust model fitting across various applications, including point estimation (Windham, 1995), structured data applications (Wang et al., 2021), clustering analysis via mean shift (Cheng, 1995), and the *γ*-estimator family (Fujisawa and Eguchi, 2008; Chen et al., 2014).

Under standard independent and identically distributed (i.i.d.) models with homoscedastic variance, the minimum density power divergence estimate (MDPDE) for both the Gaussian mean (Basu et al., 1998) and linear regression parameters (Ghosh and Basu, 2013) yields weighted formulas that are functionally equivalent to maximum likelihood estimates (MLE) for heteroscedastic models. Our strategy leverages this relationship by using MDPDE-derived weights to construct the covariance matrices 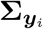 and **Λ**_*i*_ within the heteroscedastic frame-work defined in Equations (3) and (4). This approach allows us to maintain a likelihood-based hierarchy while incorporating the inherent robustness of the density power divergence method.

The density power divergence between two density functions *g*_*n*_ and *f*, where *g*_*n*_ usually refers to the empirial density function, is defined as:

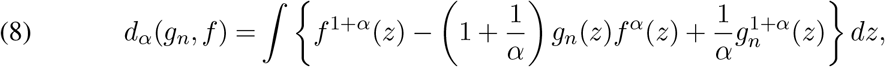

where *α >* 0 controls the trade-off between robustness and efficiency (Basu et al., 1998). Given an empirical density function *g*_*n*_ and a parametric family *{f*_***t***_, ***t*** ∈ *B}*, the MDPDE is found by minimizing the divergence: 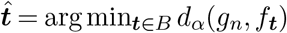.

As *α* → 0, the power divergence converges to the Kullback-Leibler divergence:

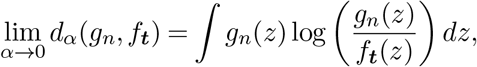

which leads to the maximum likelihood estimate (MLE). Thus, while the MDPDE reduces to the most efficient MLE at *α* = 0, increasing *α* enhances robustness against data contamination at the cost of some efficiency. It is suggested in Basu et al. (1998) to use a small value of *α*, as this balances the trade-off by introducing robustness without significantly compromising efficiency. We describe an automated cross-validation procedure for selecting this parameter in Section 2.3.

The estimating equation is derived by setting the derivative of Equation (8) to zero:

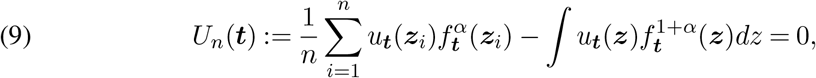

where 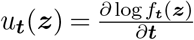 denotes the score function.

In the following sections, we describe the construction of 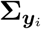 (Section 2.2.1), detail the specification of **Λ**_*i*_ (Section 2.2.2), and present the cross-validation procedure for selecting the tuning parameters, *α*_*r*_ and *α*_*g*_ (Section 2.3). Specifically, within our two-level frame-work, the parameter ***t*** from the foundational MDPDE theory refers the coefficient vector ***β*** at the atom-level regression stage and corresponds to the mean vector ***µ*** at the group-level estimation stage.

#### 2.2.1. Construction of Σ_y_ for the Linear Regression Model

Let 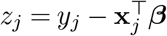 denote the residuals, and assume 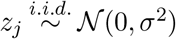 for the baseline linear regression model. The density function and the score function with respect to ***β*** are given by:

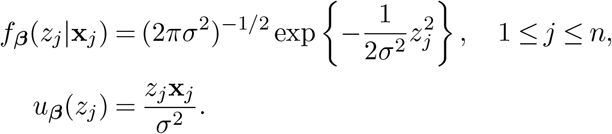

In this symmetric Gaussian case, the integral term in the estimating Equation (9) vanishes. Using the regression tuning parameter *α*_*r*_, we obtain the following estimating equation for ***β***:

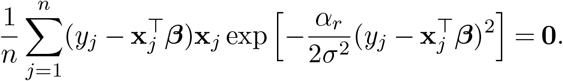

The resulting MDPDE for ***β*** can be expressed in a weighted least squares form:

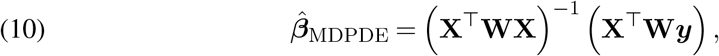

where **X** = (**x**_1_, …, **x**_*n*_)^⊤^ and **W** is a diagonal weight matrix with with the *j*^*th*^ element defined as:

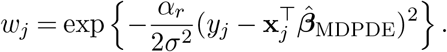

Since 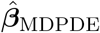 appears on both sides of Equation (10), we employ a fixed-point iterative algorithm for the solution. Using the ordinary least squares (OLS) estimate as the initial value, convergence is typically achieved in fewer than 10 iterations.

Crucially, the MDPDE for the homoscedastic model is functionally equivalent to the maximum likelihood estimate (MLE) of a heteroscedastic model where each observation (**x**_*j*_, *y*_*j*_) is assigned a weight *w*_*j*_. We thus define the equivalent heteroscedastic covariance matrix as:

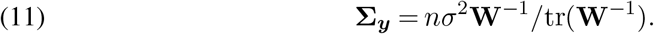

This normalization ensures that tr(**Σ**_***y***_) = *nσ*^2^, matching the trace of the homoscedastic covariance *σ*^2^**I**_*n*_. This preserves the “total information” scale, allowing for consistent variance comparisons across different atoms.

To estimate the scale parameter *σ*^2^, we use the score function 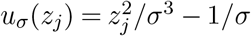 and and derive the following MDPDE for the variance:

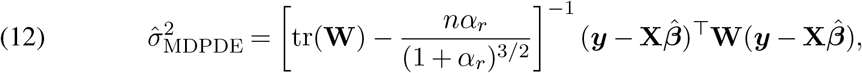

where the technical derivation is provided in A.4.1 of the Supplementary Material (Tu et al., 2026). In practice, for each atom *i* in the group, we calculate the individual covariance matrix 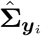 as:

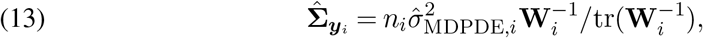

which is then substituted into the group-level estimating equations to determine the robust mean 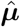.

#### 2.2.2 Construction of Λ_i_ for the Normal Distribution

To model the distribution of atomic characteristics within a category, we assume that the atom-specific parameters ***β***_*i*_ are independent realizations from a group-level distribution such that: ***β***_*i*_ ∼ *N* (***µ*, Λ**) for 1 ≤ *i* ≤ *I*, the score function and density power are given by *u*_***µ***_(***β***_*i*_) = **Λ**^−1^(***β***_*i*_ − ***µ***) and:

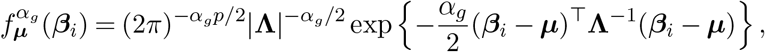

where the tuning parameter *α*_*g*_ specifically controls the robustness of the group-level estimates. In this Gaussian case, the integral term in Equation (9) vanishes, and the estimating equation simplifies to:

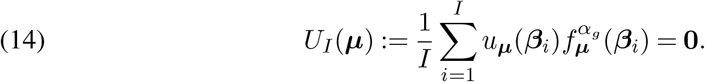

The resulting MDPDE for the group mean is a weighted average:

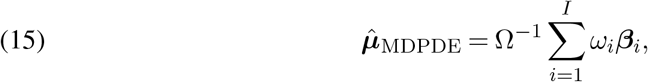

where 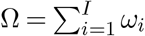 and the weights are defined as:

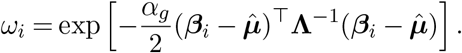

Since 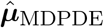 is present within the weight *ω*_*i*_, we utilize a fixed-point iterative algorithm to obtain the solution. Initializing with the sample mean, convergence is typically achieved within 10 iterations.

Following the analogy of the voxel-level regression, we interpret 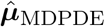 as the MLE of a heteroscedastic model where atom-specific parameters follow:

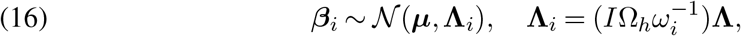

where 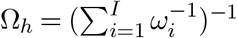 is the harmonic mean of the weights. Under this formulation, the sum of the atom-specific covariances remains *I***Λ**, aligning with the total covariance of the baseline i.i.d. model. Here, *ω*_*i*_ effectively represents the relative contribution of the *i*^*th*^ atom to the group mean, allowing the hierarchical structure to accommodate members that deviate from the population consensus.

While Basu et al. Basu et al. (1998) established asymptotic properties for the MDPDE under general frameworks, the specificity of the Normal model allows us to directly establish consistency; we show that 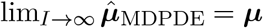 in A.4.2 of the Supplementary Material (Tu et al., 2026).

Furthermore, while existing MDPDE derivations often assume scalar or diagonal covariance structures, we provide an explicit estimator for a fully unstructured covariance matrix **Λ**:

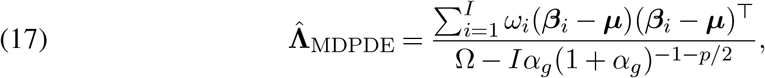

where *p* denotes the dimensionality of ***β***_*i*_ (with *p* = 2 in this study, representing log-amplitude and log-width). This generalization enables a rigorous treatment of the correlated atomic features inherent in cryo-EM data. The full technical derivation is detailed in A.4.3 of the Supplementary Material (Tu et al., 2026).

In implementation, for each atom *i*, we calculate the individual covariance 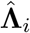 as:

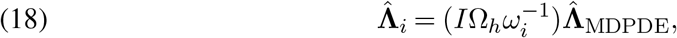

which is then substituted back into the hierarchical framework to finalize the robust group estimate 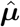.

#### 2.2.3. Theoretical Robustness and Influence Properties

To establish the theoretical grounding of the robust hierarchical linear (RHL) framework, we evaluate the robustness of the group-level mean estimate *µ* using the influence function (IF). As detailed in A.5 of the Supplementary Material (Tu et al., 2026), we specifically contrast the IF of the standard Maximum Likelihood Estimate (MLE) with that of the Minimum Density Power Divergence Estimator (MDPDE) for the population-level parameter.

While the MLE exhibits a linear and unbounded influence function, the MDPDE possesses a redescending property that ensures highly deviated data points have a bounded and ultimately negligible impact on the group-level estimates. This mathematical structure leads to a “robustness paradox” where neutralized contamination is effectively ignored by the estimator; consequently, we see the possibility to leverage the relative weight profile Ω^−1^*ω*_*i*_ as our primary diagnostic tool for identifying structural heterogeneity and structural model misspecification.

### 2.3. Selection of Tuning Parameters

The tuning parameters *α*_*r*_ (regression-level) and *α*_*g*_ (group-level) in the MDPDE framework control the fundamental trade-off between robustness and statistical efficiency. At a value of zero, the MDPDE reduces to the maximum likelihood estimator, achieving optimal efficiency under a perfectly specified model. However, in the presence of data contamination, positive values for these tuning parameters enhance robustness by down-weighting observations that deviate from the assumed Gaussian signal profile.

To identify the optimal values for *α*_*r*_ and *α*_*g*_, we employ a cross-validation strategy. The dataset is repeatedly and randomly partitioned into training and testing subsets. For each candidate tuning parameter, atom-specific and group-level estimates are derived from the training set and evaluated for their predictive stability on the testing set. The optimal parameters are those that minimize the discrepancy between training and testing estimates, thereby identifying the regime that maximizes both robustness to noise and generalizability across the protein structure. The specific implementation details for this selection process are summarized in Algorithms 4 and 5 in Section 2.4.

### 2.4. Algorithms and Codes

This section summarizes the computational procedures used to implement the proposed framework. The estimation pipeline consists of five algorithms with sequential dependencies.

Algorithm 1 computes the atom-level MDPDE estimates 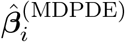 and the corresponding covariance estimates 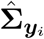. These quantities serve as inputs for subsequent stages. Algorithm 2 uses the atom-level estimates to obtain the group-level MDPDE estimate of the population mean ***µ*** and the associated covariance matrices **Λ**_*i*_. Given these estimates, Algorithm 3 evaluates the marginal likelihood and produces the empirical Bayes estimate 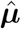.

Algorithms 4 and 5 select the tuning parameters that control robustness. Algorithm 4 determines the regression-stage tuning parameter *α*_*r*_, which is used in Algorithm 1, and Algorithm 5 selects the group-level tuning parameter *α*_*g*_ for Algorithm 2. Together, these procedures define a complete estimation workflow for the hierarchical model.

The codes in running these algorithms are available in the Github: https://github.com/yslianTim/rhbm_gem.

#### Algorithm 1

MDPDE for ***β***

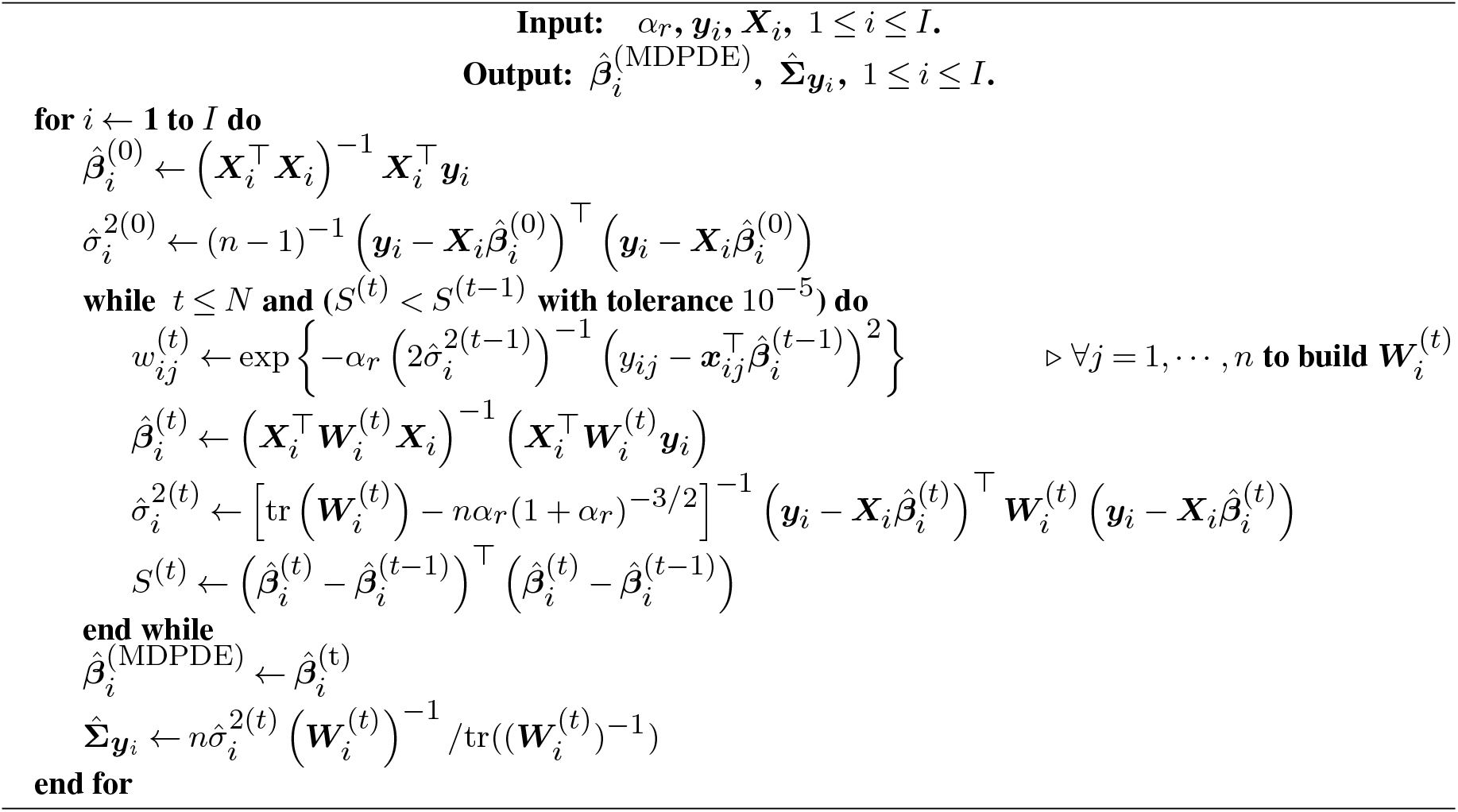

#### Algorithm 2

MDPDE for ***µ***

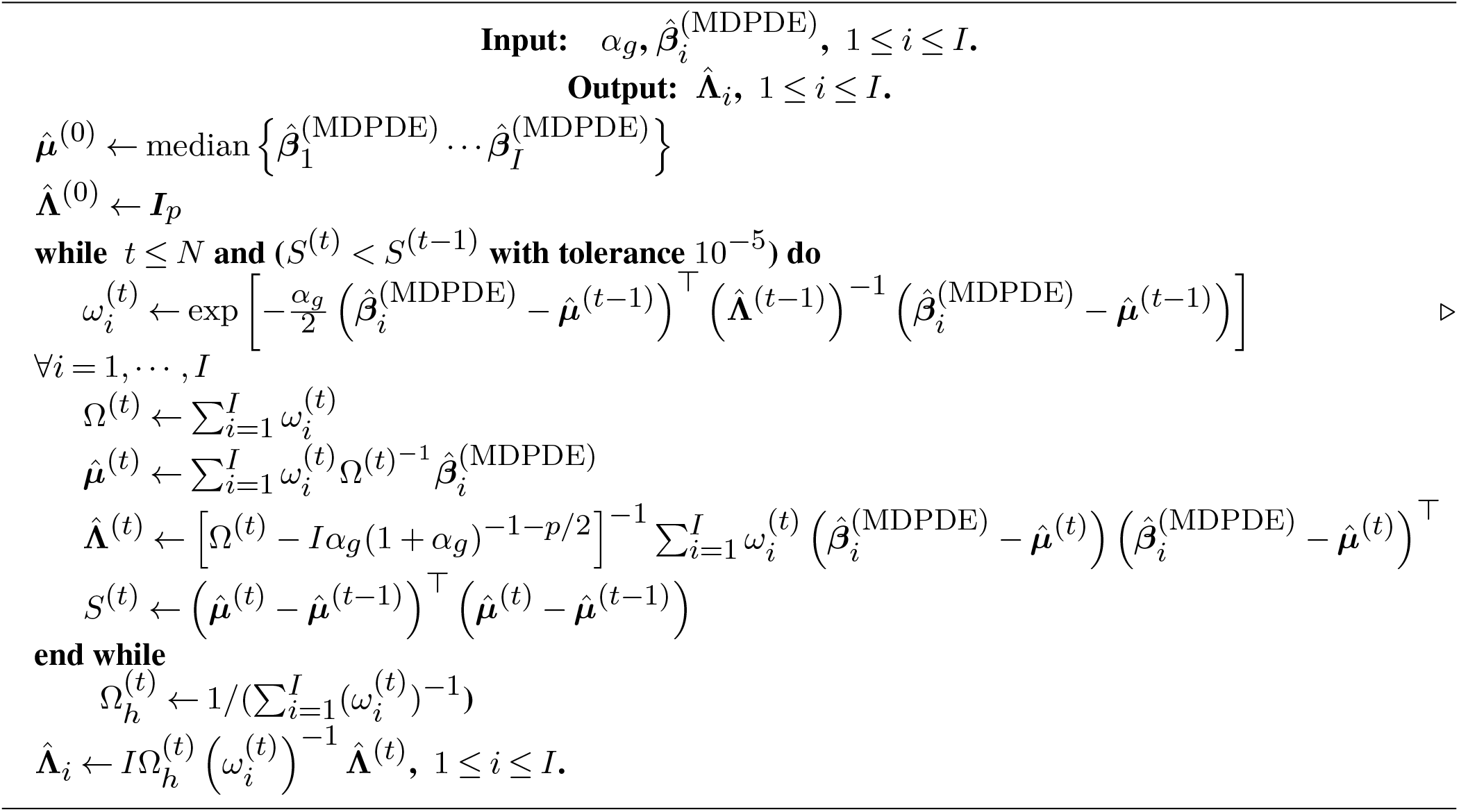

#### Algorithm 3

To Calculate the Maximum Marginal Likelihood Estimate 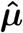

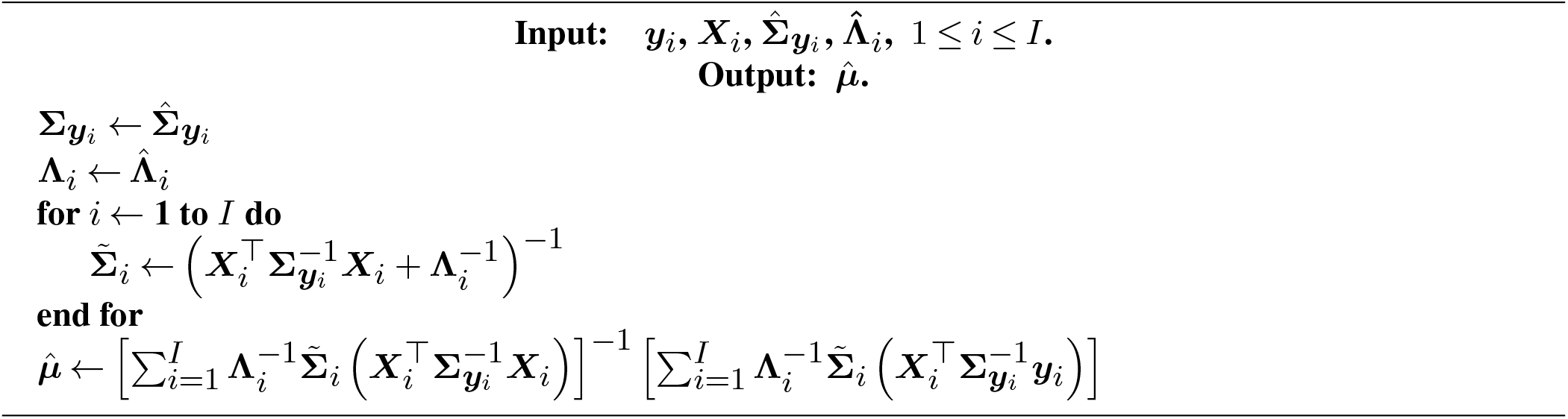

#### Algorithm 4

Cross-Validation for finding Universal *α*_*r*_

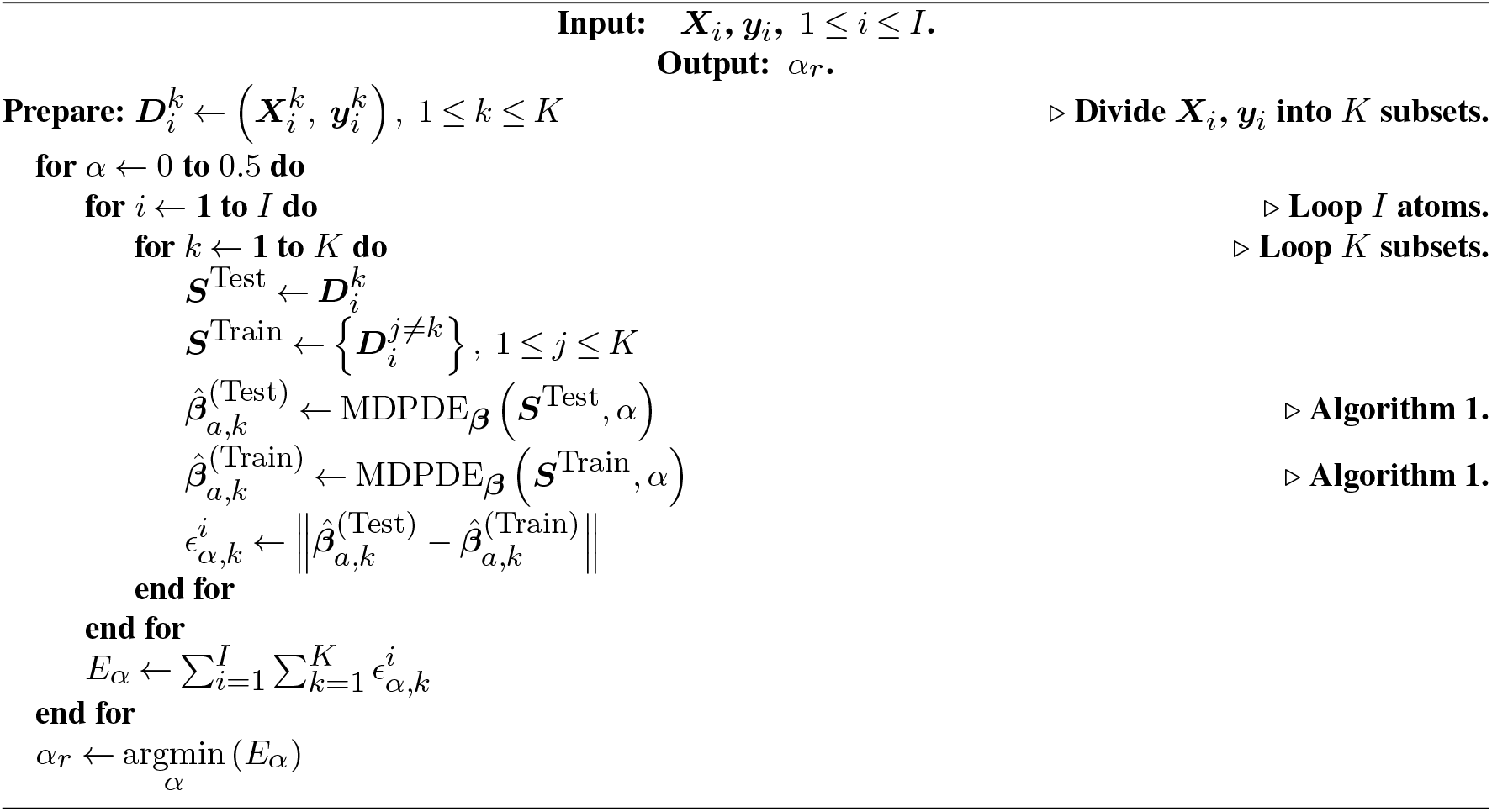

#### Algorithm 5

Cross-Validation for finding Universal *α*_*g*_

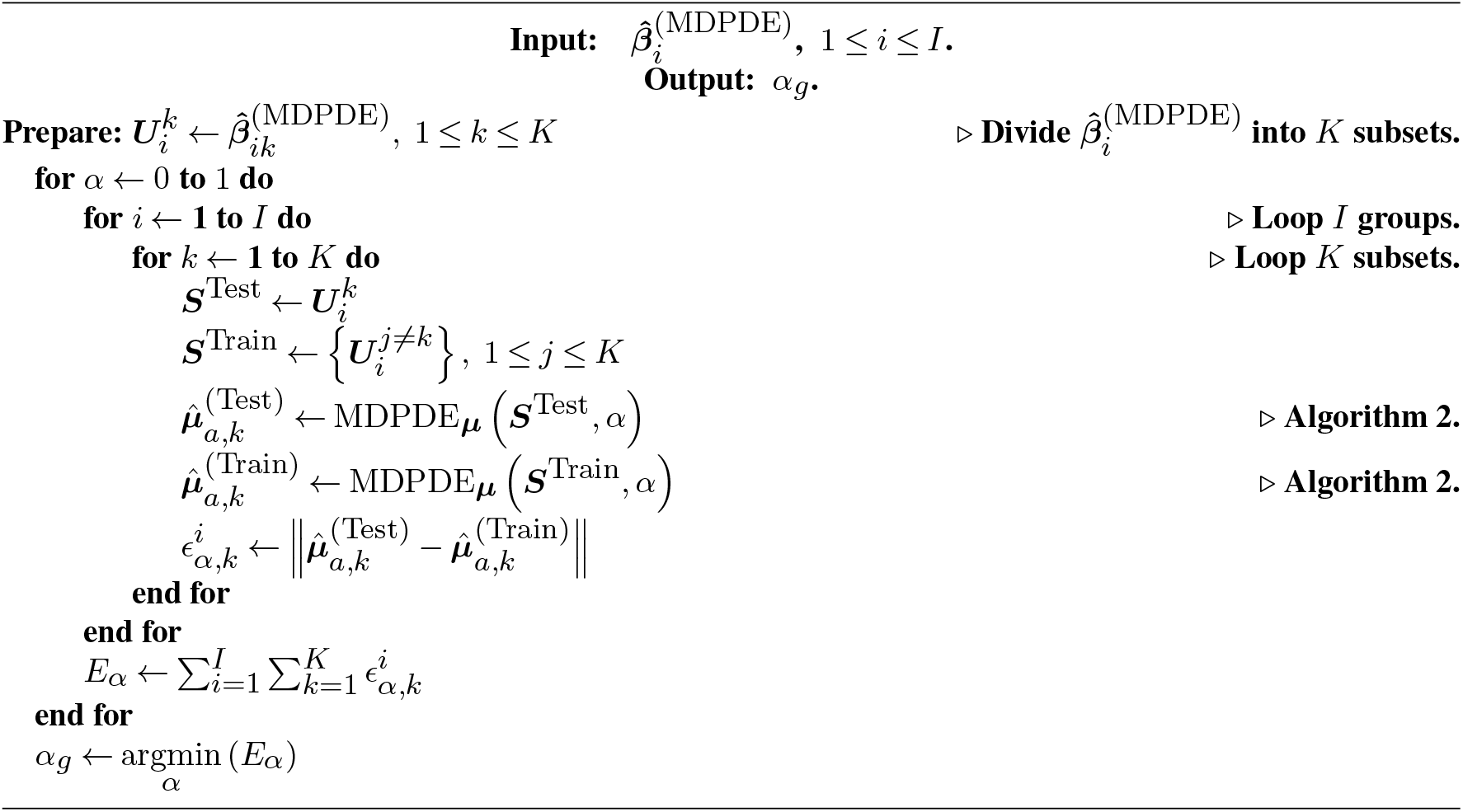

## 3. Numerical Studies

### 3.1. Simulation Study

In this simulation study, our goal is to compare the performance of robust and non-robust estimates within the hierarchical modeling framework: Equations (3) and (4), particularly under conditions involving data contamination. Notably, the estimates can be distinguished by the power parameter *α*, where a positive value refers to a robust estimate and a zero value for the MLE (non-robust). Here, the parameter *α* is of the power in Minimum Density Power Divergence Estimate (MDPDE).

In our hierarchical framework, we incorporate two levels of statistical models: the bottom level represents the regression model, and the top level integrates group-specific information. We introduce *α*_*r*_ to denote the power parameter at the regression level and *α*_*g*_ to signify the power parameter at the group integration level, where *α*_*r*_ = 0 refers to the ordinary least squares estimate and *α*_*g*_ = 0 refers to the mean estimate. The bias ratio (normalized by the true parameter) of the estimate serves as the primary metric for comparing performance, to measure the impact from the data contamination.

We first consider a linear regression setting and compare ordinary least squares (OLS; *α*_*r*_ = 0), the MDPDE with a fixed tuning parameter (*α*_*r*_ = 0.1), and the MDPDE with the tuning parameter selected by Algorithm 4. The data are generated from a homoscedastic linear regression model

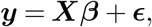

with specified proportions of data contamination. Here, the parameter vector ***β*** is set to be

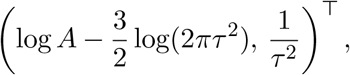

where the designed matrix ***X*** = (***x***_1_, …, ***x***_*n*_)^⊤^ is constructed with each data point ***x***_*j*_ given by 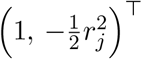, and parameters (*A, τ*) = (1.0, 0.5). The error terms are sampled from the Normal distribution 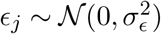. To investigate the effect of varying error levels, we tested three values of *σ*_*ϵ*_: 0.1*D*_max_, 0.2*D*_max_ and 0.3*D*_max_, where *D*_max_ = 0.508 is the peak of the 3D Gaussian density function, given by *Aϕ*(0; *τ* ^2^). Data contamination is generated under the same model specification, but with a modified parameter 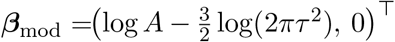, which eliminates the relationship between the predictors and response. Data examples for the three error intensities with 10% data contamination are illustrated in Figure 2.

**FIG 2.**
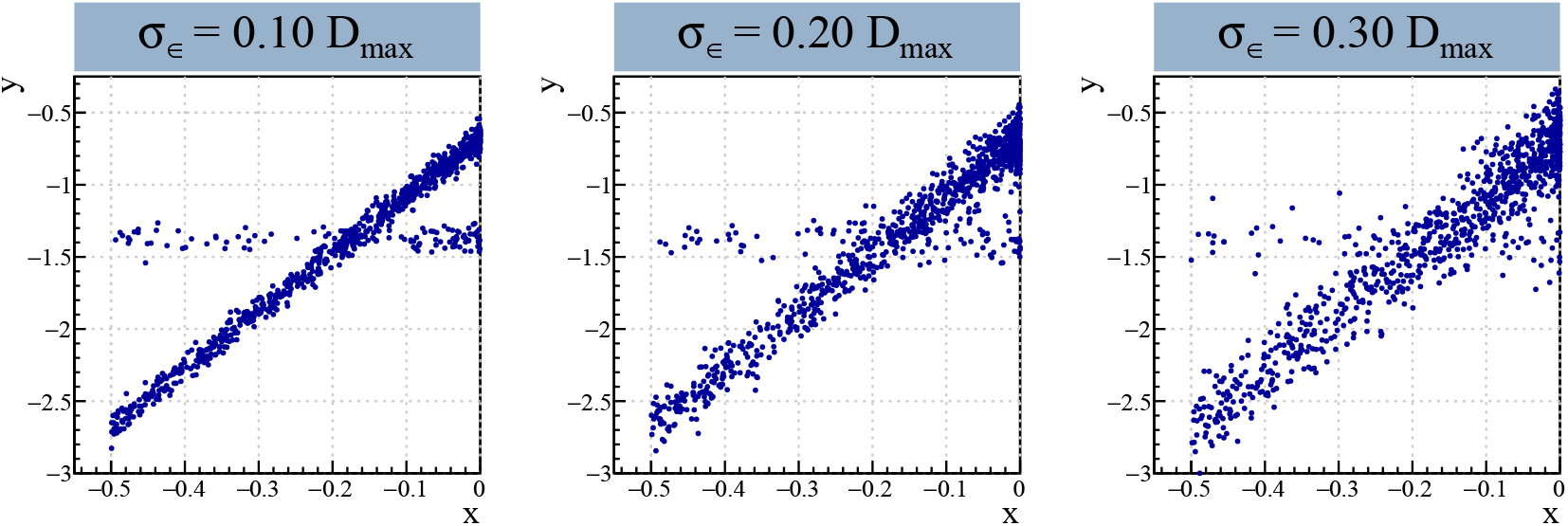
Illustration of the Simulation Data: We generate 900 data points following the model **y** = **Xβ** +**ϵ**, where 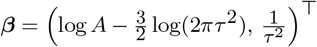, and (A, τ) = (1.0, 0.5). **ϵ** is sampled from the Normal distribution 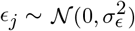 with 3 intensities for σϵ: 0.1Dmax, 0.2Dmax and 0.3Dmax, where Dmax = 0.508 = Aϕ(0; τ^2^). Additionally, 100 data points (10% of the dataset) are generated as data contamination using the same model structure but with a modified parameter: 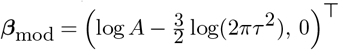, which removes the relationship between predictors and the response.

In each simulation setting, we generate a sample of size 1,000 and repeat the experiment 100 times to evaluate the bias and standard deviation of the estimates under contamination ratios ranging from 0 to 20%. For each contamination level, we compare the performance of three estimators: (i) ordinary least squares (OLS), corresponding to *α*_*r*_ = 0; (ii) the MDPDE with a fixed tuning parameter *α*_*r*_ = 0.1; and (iii) the MDPDE with *α*_*r*_ dynamically selected using the cross-validation procedure in Algorithm 4.

To obtain the physical signal features, we first employ the MDPDE framework described in Section 2.2.1 to estimate the regression coefficient vector 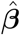. We then apply the transformation defined in Equation (2) to derive the corresponding estimates for atomic amplitude (*Â*) and width 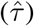.

The resulting performance is summarized in Figure 3. We report the normalized bias and variability, defined as the mean and standard deviation of the 100 values 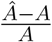 and 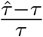 across simulation replicates.

**FIG 3.**
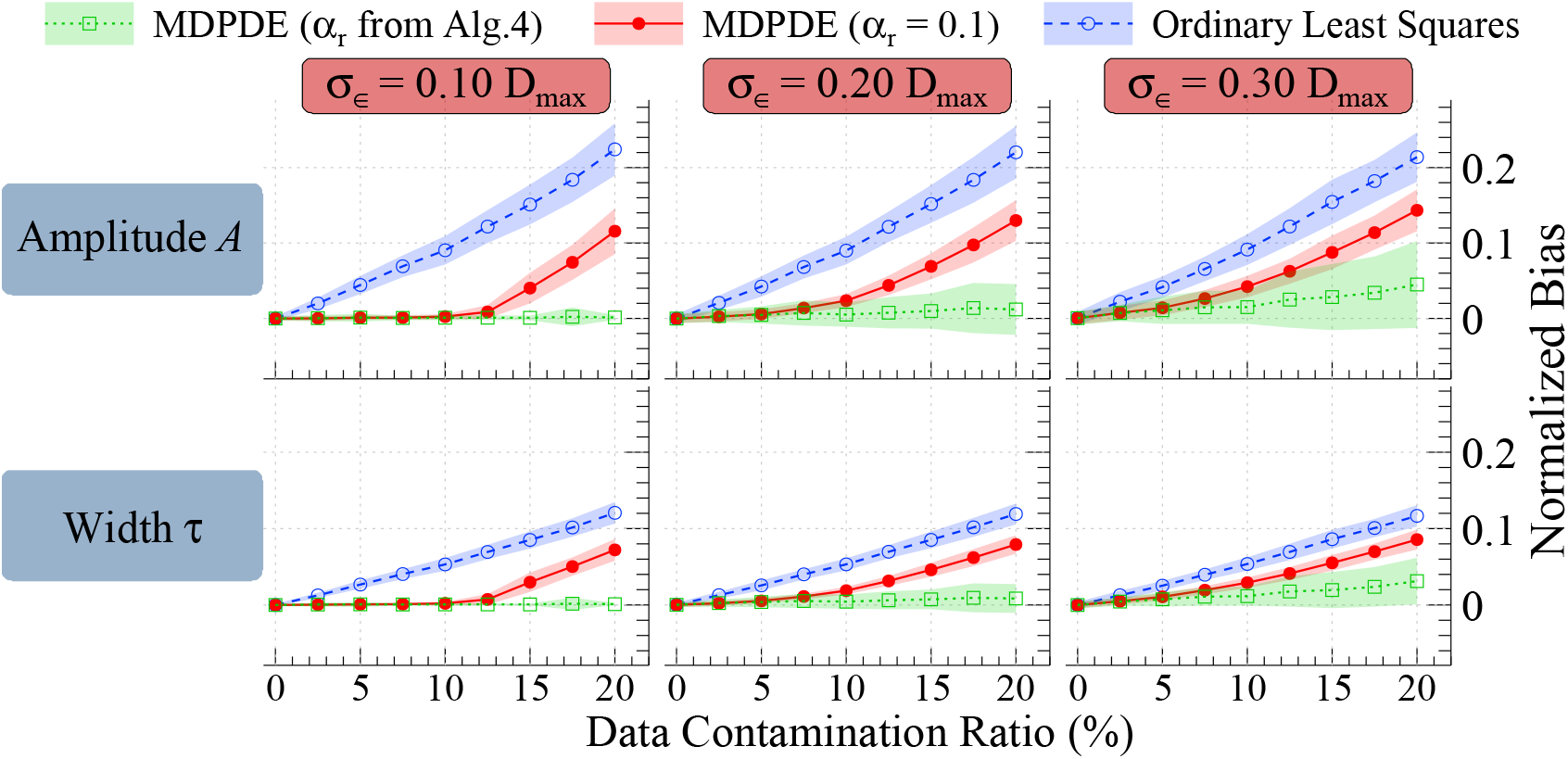
Bias comparison of OLS and MDPDE under data contamination. Using the simulated data shown in Figure 2, we evaluate estimation performance for ordinary least squares (OLS) and minimum density power divergence estimation (MDPDE). Normalized bias for the amplitude parameter (top) and width parameter (bottom) are plotted against the contamination proportion. Markers denote the mean normalized bias, and shaded vertical bands represent the corresponding standard deviations on both sides. MDPDE results are shown in open square (αr selected using Algorithm 4) and in solid circle (αr = 0.1), while OLS results are shown in open circle. The three columns correspond to three levels of error intensity. The results indicate that MDPDE yields substantially smaller bias than OLS across contamination levels, and that the cross-validated tuning parameter further reduces bias compared with the fixed choice in this simulation.

Figure 3 shows that MDPDE outperforms OLS in the presence of contamination, while performing comparably to OLS when no contamination is present. For a contamination ratio of 10%, MDPDE with a small tuning parameter (*α*_*r*_ = 0.1) effectively mitigates the impact of contaminated observations. However, as the contamination ratio increases, MDPDE with the fixed choice *α*_*r*_ = 0.1 becomes less resistant to contamination. In contrast, MDPDE with *α*_*r*_ selected using Algorithm 4 continues to perform well across contamination levels.

To further illustrate the effect of the tuning parameter, we present additional bias and variability comparisons for two fixed tuning parameters in Figure 2 of the Supplementary Material (Tu et al., 2026), highlighting the trade-off between robustness (bias ratio) and efficiency (standard deviation).

Secondly, we evaluate the estimation performance of the hierarchical model when data contamination occurs at the atom level. Specifically, we compare three estimators for the population parameters: (i) the non-robust estimator (MLE, *α*_*g*_ = 0); (ii) the MDPDE with a fixed tuning parameter (*α*_*g*_ = 0.2); and (iii) the MDPDE with *α*_*g*_ dynamically selected using Algorithm 5. For each replicate, we first compute the group-level mean 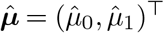 from Equation (15) using the iterative procedure in Algorithm 2. The corresponding group-level amplitude (*Â*) and width 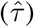 are then obtained via the following transformation:

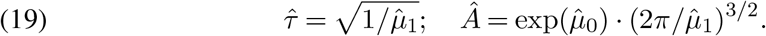

To evaluate the model’s performance under conditions mimicking real-data applications, we simulate distributions for the atom-specific physical parameters *A*_*i*_ (amplitude) and *τ*_*i*_ (width). In the uncontaminated (null) setting, we assume:

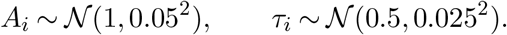

Level-2 (atom-level) contamination is introduced by applying alternative distributions to a randomly selected subset of indices *i* ∈ {1, …, *I}* corresponding to a specified contamination proportion. We investigate two distinct contamination regimes:

#### Amplitude Contamination

Contaminated atoms follow *A*_*i*_ ∼ *N* (1.5, 0.05^2^) while the distribution for *τ*_*i*_ remains unchanged.

#### Width Contamination

Contaminated atoms follow *τ*_*i*_ ∼ *N* (1.00, 0.025^2^) while the distribution for *A*_*i*_ remains unchanged.

For each experimental configuration, we simulate I=100 groups across 100 replicates to compute the bias and standard deviation of the estimates, with contamination ratios ranging from 0% to 20%.

The resulting performance is summarized in Figure 4. We report three estimators: the non-robust estimator (*α*_*g*_ = 0), the robust estimator with fixed *α*_*g*_ = 0.2, and the estimator with *α*_*g*_ selected using Algorithm 5. The normalized bias and variability are defined as the mean and standard deviation of the 100 values 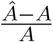 and 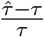 across simulation replicates.

**FIG 4.**
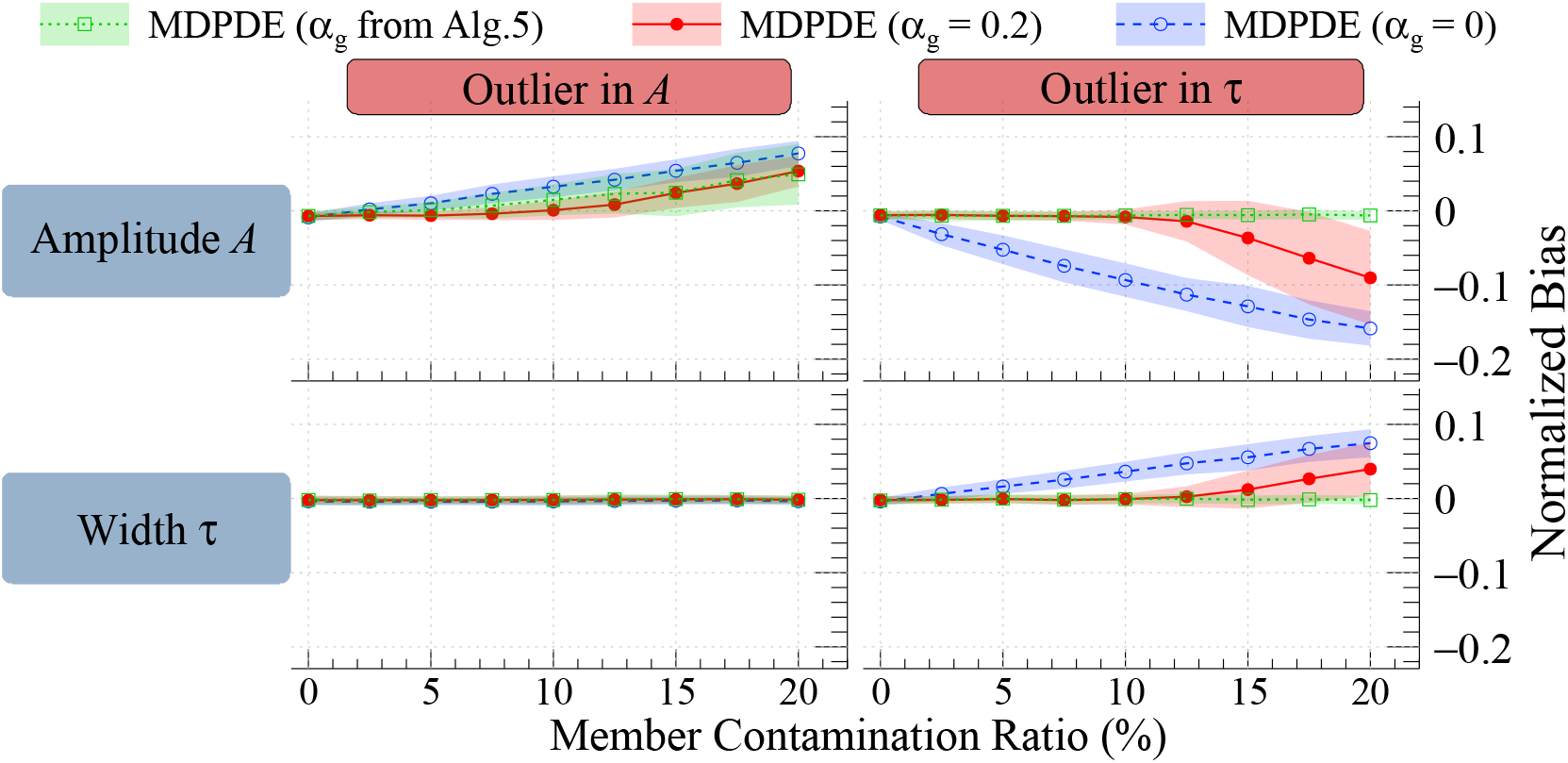
Bias comparison of group-level parameter estimates under data contamination. Normalized bias for the amplitude parameter (top) and width parameter (bottom) are plotted against the contamination proportion. Markers denote the mean normalized bias, and shaded vertical bands represent the corresponding standard deviations on both sides. MDPDE results are shown in open square (αg selected using Algorithm 5) and in solid circle (αg = 0.2), while the mean estimator (MLE, αg = 0) is shown in open circle. The left column corresponds to contamination introduced in amplitude, and the right column corresponds to contamination introduced in width. The coincidence of the three markers in the lower-left panel indicates that contamination in amplitude has negligible impact on width estimation for both the robust and non-robust estimators in this setting.

Figure 4 shows that the robust estimator with *α*_*g*_ = 0.2 outperforms the non-robust estimator (*α*_*g*_ = 0) in the presence of contamination, while performing comparably when no contamination is present. For a contamination ratio of 10%, the robust estimator with the fixed tuning parameter *α*_*g*_ = 0.2 effectively mitigates the impact of contaminated observations. However, as the contamination ratio increases, its resistance to contamination gradually decreases. In contrast, the estimator with *α*_*g*_ selected using Algorithm 5 performs comparably for amplitude estimation and more consistently for width estimation across contamination levels.

To further illustrate the influence of the tuning parameter, we present additional bias and variability comparisons for two fixed tuning parameters in Figure 3 of the Supplementary Material (Tu et al., 2026), highlighting the trade-off between robustness (bias ratio) and efficiency (standard deviation).

An interesting observation from Figure 4 is that when data contamination affects only the amplitude, the bias appears only in the amplitude estimate, indicating that amplitude data contamination does not impact width estimation for either robust or non-robust estimates. Conversely, when data contamination influences the width parameter, bias emerges in the estimates for both the width and amplitude parameters, as shown by the markers across both panels.

#### 3.2. Real Data Analysis

#### 3.2.1. Three Cryo-EM Maps Analysis: Resolution-Dependent Behavior of Width Estimates

We apply our proposed framework to three experimental cryo-EM maps with reported Fourier Shell Correlation (FSC) resolutions of 1.25 Å (Yip et al., 2020), 2.20 Å (Cianfrocco et al., 2018), and 3.23 Å (Nirwal et al., 2025). For each of the three backbone atom types across the 20 standard amino acids, we compute the group-level estimator 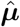 using the robust procedure defined in Equation (7). In this implementation, the regression-level tuning parameter *α*_*r*_ is dynamically selected via Algorithm 4, and the group-level parameter *α*_*g*_ is selected via Algorithm 5.

The resulting group-level amplitude (*Â*) and width 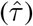 estimators are subsequently derived through the parameter transformation in Equation (19). The estimated signal features for the three backbone atoms across all 20 amino acid types for each map are summarized in Figure 5.

**FIG 5.**
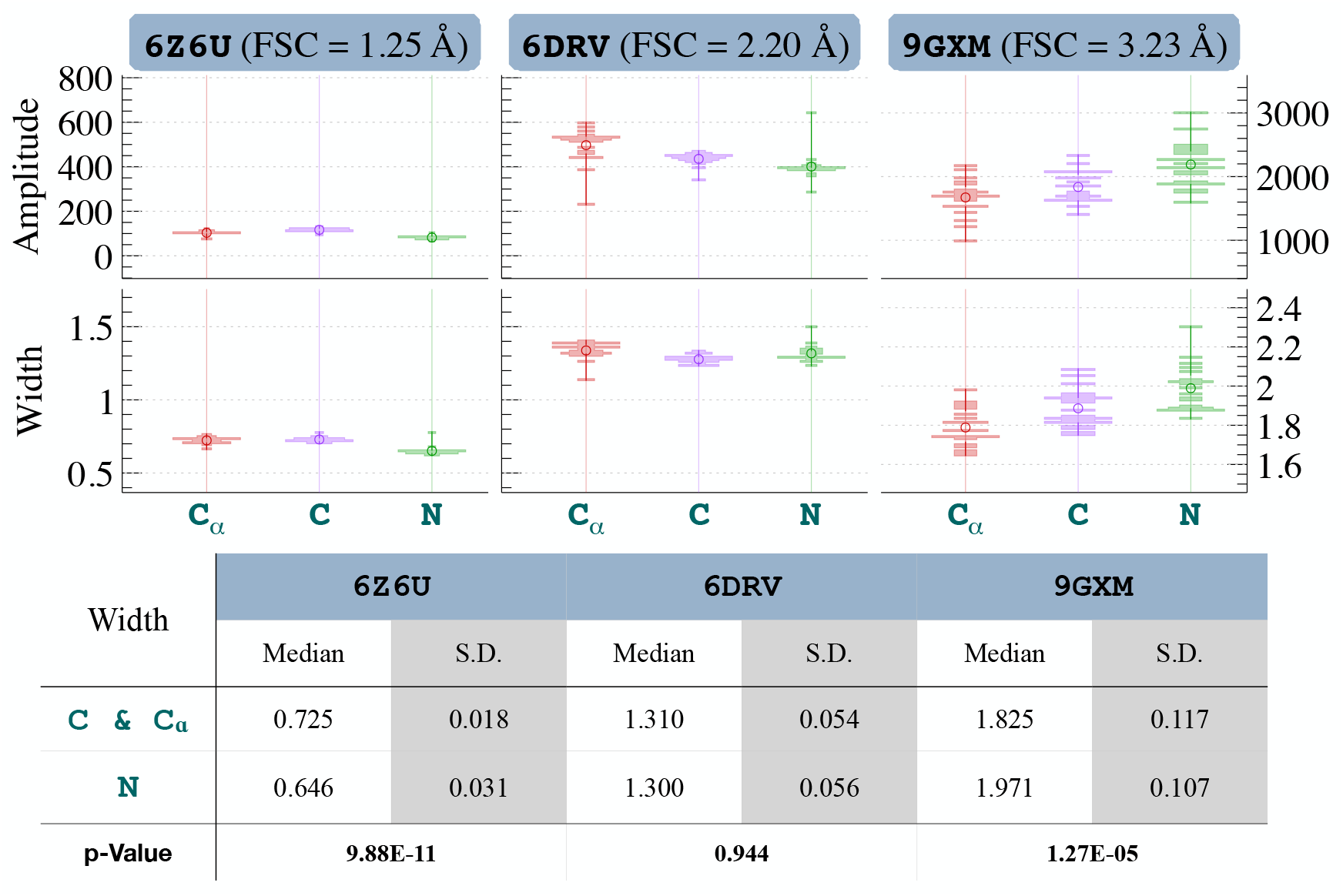
Estimates for Three Cryo-EM Maps. The figure displays estimated Gaussian amplitude and width parameters for three backbone atom types across three cryo-EM maps Yip et al. (2020); Cianfrocco et al. (2018); Nirwal et al. (2025). For each map and atom type, 20 estimates corresponding to the 20 amino acid types are summarized. Medians are shown as circles, and the width of each band is proportional to the frequency of observations at the corresponding values. Amplitude estimates are not directly comparable across maps due to mapspecific scaling; in particular, 6Z6U and 6DRV are on comparable scales, whereas 9GXM is not. In contrast, width estimates show a clear association with the reported FSC resolutions. The accompanying table reports the median and standard deviation of the 20 width estimates for carbon (combining C and Cα) and N, along with p-values from the Wilcoxon rank-sum test assessing differences between the two atom types within each map.

Since amplitude estimates are highly sensitive to map-specific scaling and experimental conditions, direct cross-map comparisons of raw intensities are not readily interpretable. In contrast, the width estimates 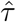, which quantify signal dispersion at the atomic level, demonstrate a clear association with the reported FSC resolutions. The table in Figure 5 summarizes the median and standard deviation of the width estimates across the 20 amino acids for each map, where the carbonyl carbon (C) and *α*-carbon (*C*_*α*_) are analyzed as a combined group.

Across the three maps, the distribution of width estimates reflects the underlying structural resolution: lower-resolution data are characterized by greater variability and increased dispersion. Specifically, the width estimates for the 1.25 Å map (6Z6U) are tightly concentrated, whereas the 3.23 Å map (9GXM) exhibits a heavy-tailed distribution, and the 2.20 Å map (6DRV) shows noticeable skewness. Because the distributions of these width estimates deviate from normality, we utilize the median as a robust summary statistic to compare results across different atom types. Notably, these estimates are derived independently for each amino acid category; our framework does not impose a prior distribution model on these parameters.

To determine if width estimates systematically differ between chemical species, we applied the Wilcoxon rank-sum test to compare the distributions of the Carbon group (C and *C*_*α*_) and the Nitrogen group. For the atomic resolution 1.25 Å map, the test yielded a highly significant p-value (9.88 *×* 10^−11^), indicating a distinct signal profile for Nitrogen relative to Carbon. In contrast, this difference was not statistically significant for the 2.20 Å map. While the test reached significance for the 3.23 Å map, the relative ranking of the atom types became unstable, likely reflecting the reduced spatial precision inherent in lower-resolution data. These results suggest that the detectability of atom-specific signal features is strongly resolution-dependent, with ultra-high-resolution maps providing the most reliable statistical evidence for these subtle physical differences.

To further illustrate the capacity of our framework to extract nuanced atomic-level features, we next present a detailed analysis of the 1.25 Å apoferritin map (EMD-11103 / PDB 6Z6U).

#### 3.2.2. Zoom-in Analysis for 6Z6U

PDB entry 6Z6U, associated with EMDB-11103, represents the 1.25 Å resolution cryo-EM benchmark protein, apoferritin (Yip et al., 2020). This assembly comprises 4,128 amino acids arranged with 24-fold symmetry; consequently, when treating all amino acid categories collectively, the structure yields I=172 independent observations for our hierarchical model.

Figure 6 presents the extracted map values categorized by the three backbone atom types: *C*_*α*_, *C*, and *N* . For visualization, the three-dimensional voxel data for each atom are summarized as one-dimensional radial profiles by calculating the median intensity across grid points equidistant from the atomic center. These empirical profiles are shown as solid blue lines (representing the 172 atoms across the 20 amino acid types). The red dashed curves represent the fitted Gaussian signals obtained from the MDPDE-based hierarchical estimator, which utilizes all three-dimensional voxel observations with dynamically selected tuning parameters. As shown in Figure 6, the Gaussian density function—parameterized by amplitude and width—provides a robust and informative approximation of the local atomic signal.

**FIG 6.**
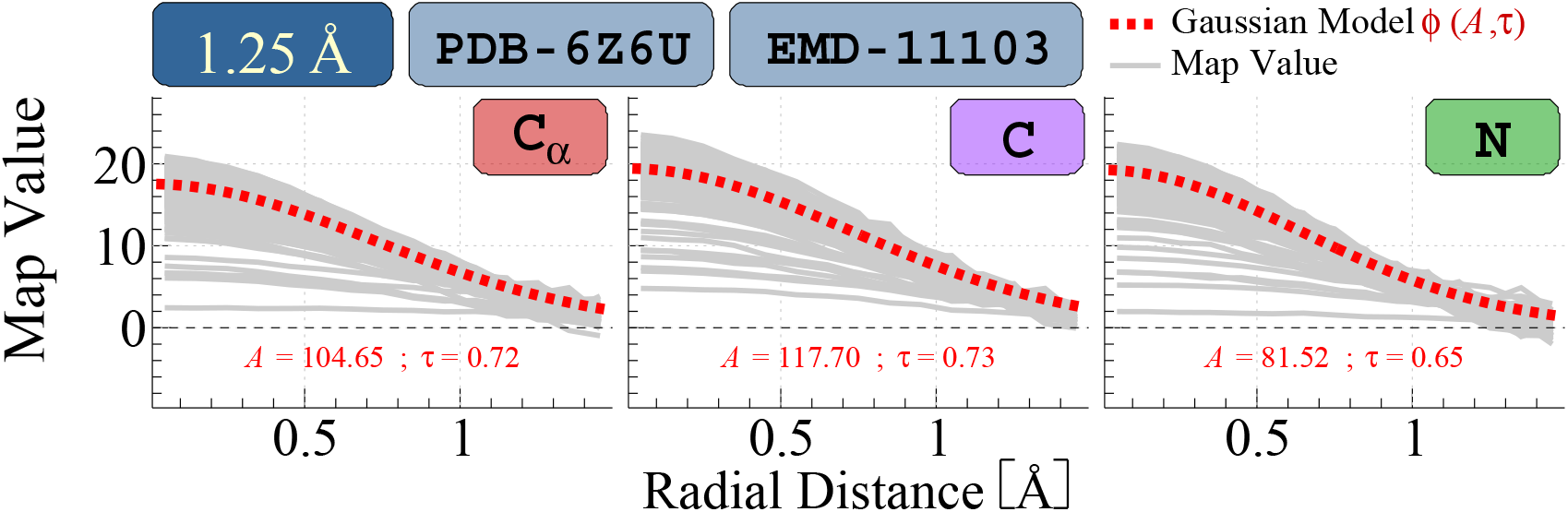
One-Dimensional Radial Profiles for Backbone Atoms (EMD-11103/PDB-6Z6U). To visualize the atomic signal features, the three-dimensional voxel intensities are summarized as one-dimensional radial profiles by calculating the median map intensity across grid points equidistant from each atomic center. These empirical profiles (solid gray lines) represent 172 independent atom-specific observations covering all 20 types of amino acids, categorized by the three backbone types: Cα, C, and N. The red dashed curves denote the fitted Gaussian signals derived from the MDPDE-based hierarchical estimator, which incorporates all underlying three-dimensional voxel observations using dynamically selected tuning parameters (αr, αg). The high degree of alignment between the empirical medians and the fitted curves demonstrates the adequacy of the Gaussian approximation for high-resolution atomic signals.

An analogous analysis, categorized by the 20 standard amino acid types, is presented in Figure 4 of the Supplementary Material (Tu et al., 2026). To further demonstrate the robustness of our framework, we present results for the *C*_*α*_ atoms of three representative residues: ILE, GLY, and SER. Figure 7 displays the corresponding Gaussian signal fits. The empirical radial profiles (solid blue) are compared against the robust MDPDE-based estimator (red dashed) and the non-robust estimator (*α*_*r*_ = *α*_*g*_ = 0; blue dotted).

**FIG 7.**
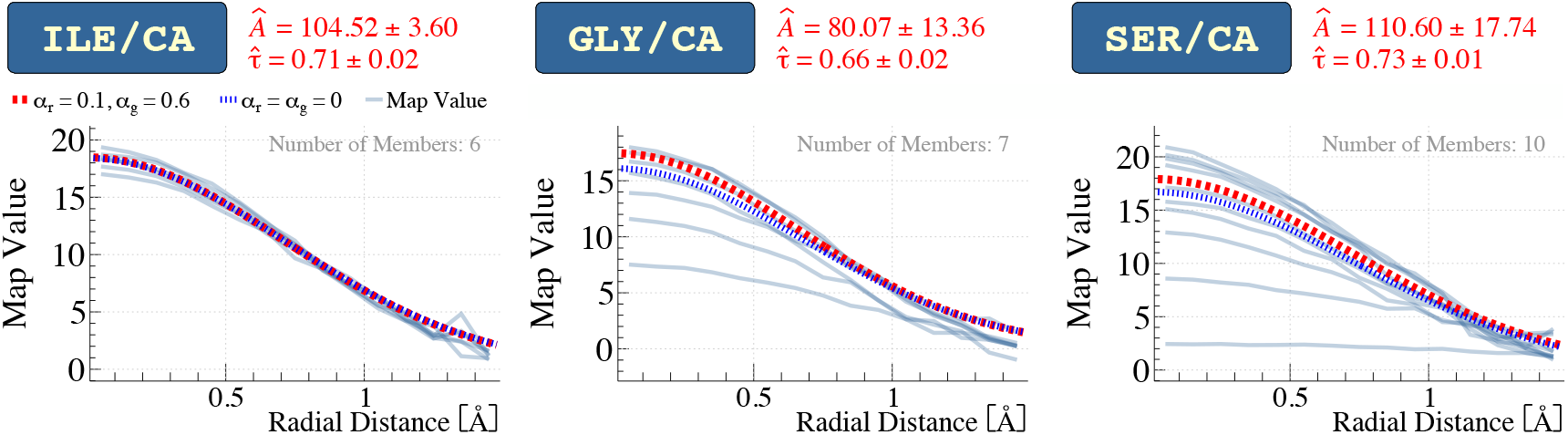
Comparison of robust and non-robust Gaussian fits for *Cα* atoms in three residue types: ILE, GLY, and SER (EMD-11103 / PDB-6Z6U). For each residue type, three-dimensional voxel intensities around individual Cα atoms are summarized as one-dimensional radial profiles (solid blue curves), obtained by taking the median map intensity across grid points equidistant from each atomic center. The red dashed curves represent robust group-level estimates obtained with data-adaptive tuning parameters (αr, αg), while the blue dotted curves correspond to non-robust estimates (αr = αg = 0). The results show that the robust estimator better preserves the dominant signal pattern in the presence of heterogeneous atom-specific observations, particularly for GLY and SER.

The amino acid ILE illustrates a homogeneous case where the atom-specific profiles are highly consistent, resulting in nearly identical robust and non-robust estimates. In contrast, GLY and SER exhibit significant heterogeneity, with several atomic profiles deviating substantially from the population consensus. In these instances, the robust estimator remains aligned with the dominant signal pattern, whereas the non-robust estimator is pulled downward by the outlier curves. This comparison highlights the advantage of the proposed frame-work in recovering reliable group-level parameters in the presence of heterogeneous atom-level observations.

To further evaluate the diagnostic utility of our atom-specific width estimates, we conducted a sensitivity analysis of the Q-score using the backbone atoms of PDB 6Z6U. We compared the Q-scores obtained using the default width parameter (0.6) against our RHL-derived estimates, categorized by atom type across three plots. Figure 8 shows that using the RHL-derived estimates presents an overall better metric; however, the improvement is only marginal. We further present the scores for two additional fixed width values, 0.5 and 0.7, as summarized in the accompanying table for four representative *C*_*α*_ atoms. The downstream validation yielded only marginal improvements in correlation that were not statistically distinguishable from the default results.

**FIG 8.**
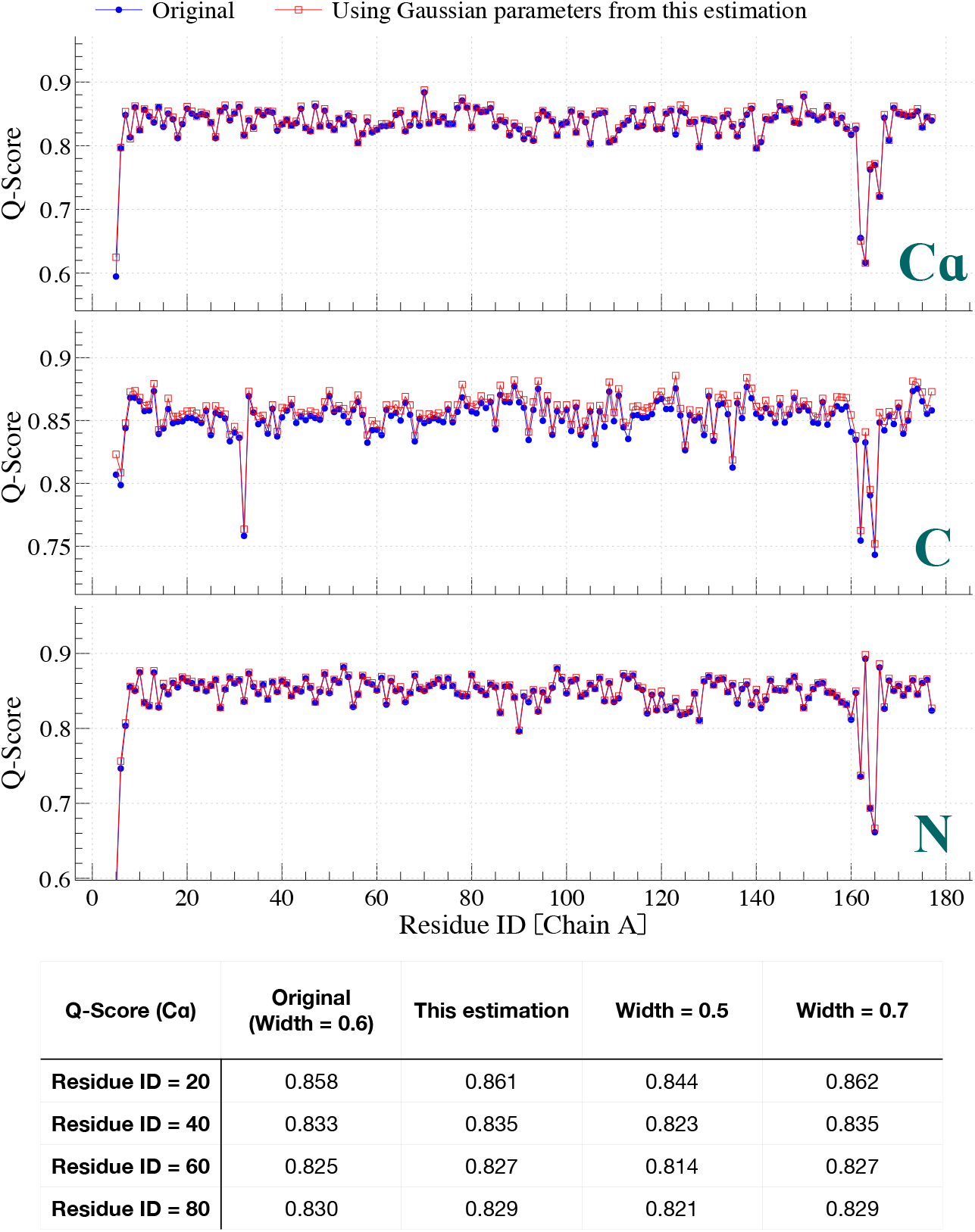
Sensitivity analysis of Q-score to Gaussian width parameters. (Top Panel): Comparison of Q-score profiles for backbone atoms (C, Cα, N) using the default width (τ=0.6) and the atom-specific parameters derived from our RHL estimation. The results across Residue IDs for Chain A show that the scores by RHL derived estimates produce better score but with only marginal improvement. (Bottom Table): A detailed sensitivity test for four representative Cα atoms (Residue IDs 20, 40, 60, and 80). The table compares Q-scores calculated using the original width, our RHL estimates, and two additional fixed widths (τ=0.5 and τ=0.7). The sampling scheme used to obtain map values for Q-score calculation is described in Section A.2 of the Supplementary Material (Tu et al., 2026). The marginal variations observed—typically within 0.01 units—demonstrate that the Pearson correlation utilized by the Q-score is relatively insensitive to the precise calibration of Gaussian width parameters. This confirms that while the RHL model achieves a statistically superior fit to the local 3D density, global metrics lack the resolution to distinguish the fine-grained, atom-specific properties or structural model misspecifications identified by our robust framework.

This observation underscores a fundamental limitation of global similarity indices, such as the Pearson correlation utilized in Q-score calculations: they lack the sensitivity required to resolve the fine-grained, atom-specific variations captured by the RHL model. While our estimated parameters provide a statistically more robust fit to the local reconstruction data, the relative invariance of the Q-score across width variations between 0.5 and 0.7 confirms that such metrics prioritize global signal trends over local structural nuances. Consequently, the RHL framework may serve as a complementary diagnostic tool, offering a layer of quantitative structural assessment that traditional correlation-based metrics are mathematically unable to resolve.

## 4. Discussion

The Robust Hierarchical Linear (RHL) framework introduced in this study provides a statistically grounded approach to quantifying atomic signals in cryoelectron microscopy maps. By moving beyond the limitations of traditional maximum likelihood estimation, our method allows for the robust detection of atom-specific features—specifically amplitude and width—while effectively neutralizing the impact of data contamination. A core methodological contribution of this work is the derivation of an explicit estimator for a fully unstructured, *p*-dimensional covariance matrix Λ. Unlike most existing MDPDE applications restricted to diagonal forms, our mathematical extension allows the RHL model to rigorously account for the inherent physical correlations between atomic parameters, such as the relationship between signal amplitude and width.

### 4.1. Methodological Constraints and Model Dependency

The efficacy of the RHL framework is fundamentally contingent upon two primary factors: the accuracy of the underlying atomic model used to anchor atomic positions, and the assumption that data contamination—or “deviated data”—constitutes a limited proportion of the observations, allowing the MDPDE to recover a representative group-level signal. Because the framework relies on these coordinates to define local sampling regions, a sufficiently accurate initial coordinate set is required to produce biophysically meaningful estimates.

A natural question is how to formalize the notion of “limited deviation” within this frame-work. The use of likelihood-based weights is conceptually related to some clustering or mode-seeking algorithms, in which the convergence to the group-representative value depends on the initial estimate lying within the basin of attraction of the dominant mode (Chen et al., 2014; Cheng, 1995). Because we use the maximum likelihood estimate (MLE) as the initial value, the robustness of the MDPDE is inherently constrained by the requirement that the MLE remains representative of the primary group. If contamination is sufficiently severe to pull the MLE toward a secondary, deviant mode, the subsequent robust weighting scheme may converge to a non-representative solution. Thus, the “limited deviation” assumption implies that, while the MDPDE can effectively downweight deviant observations, the initial estimate must remain sufficiently close to the dominant group to prevent misdirection during the iterative procedure.

In instances where atomic positions are severely misspecified, the resulting parameters may lack physical interpretability or trigger the diagnostic labeling of the atom as “deviated”. Consequently, the RHL framework should be viewed as a tool for structural validation and signal quantification within the context of a given atomic model, rather than a method for ab initio coordinate determination. While our robust weighting scheme can neutralize limited data contamination, it remains dependent on the structural model to provide the physical reference necessary for interpreting fine-grained features. Notably, by embedding these robust weights directly into the hierarchical structure, the framework maintains the analytic tractability and computational efficiency characteristic of the likelihood-based framework.

### 4.2. Spatial Correlation and Signal Preservation

A critical consideration in the analysis of cryo-EM reconstructions is the presence of spatial autocorrelation among voxel intensities. To clarify the scope, the RHL analysis focuses exclusively on local voxel values within a 1.5 Å radius of the PDB-defined atomic centers, rather than analyzing the reconstructed map as a global volume. We explicitly define these localized features as the primary atomic signals. We have adopted a signal-first strategy to address this, treating local voxel dependency as an intrinsic feature of the atomic density itself. The Gaussian density function in our model is specifically designed to capture this spatial spread as reconstructed in the map. Rather than employing aggressive noise-reduction techniques that might risk “aliasing” signal features into a complex noise model, our framework preserves the dependent voxel values to accurately characterize the physical density (the “blob”) of each atom.

Furthermore, our hierarchical framework remains robust due to the physical separation of samples. For group-level estimation (***µ***), individual atoms within a specific amino acid type are physically separated by distances that far exceed the overlap region. This ensures that the observations used to estimate population-level parameters are effectively independent, fulfilling the requirements of the hierarchical stage. By employing a heteroscedastic formulation (**Λ**_*i*_) that adaptively absorbs local density fluctuations, we maximize the preservation of chemically meaningful signal anomalies—such as the observed Nitrogen Anomaly—while maintaining model interpretability.

### 4.3. Diagnostic Utility and the Robustness Paradox

Our theoretical analysis of the influence function (IF) (A.5 of the Supplementary Material (Tu et al., 2026)) reveals what we term the “robustness paradox”. The redescending property of the MDPDE ensures that highly deviated observations have a negligible impact on the final group-level estimates. While this confirms the stability of our model under contamination, it also renders the influence function a poor indicator of neutralized contamination for our estimate. We therefore recommend weight-based diagnostics, such as the relative weight profile *ω*_*i*_*/*Ω, as the primary metric for identifying structural heterogeneity. Because *ω*_*i*_ is proportional to the likelihood raised to the power *α*_*g*_, deviant observations yield lower likelihood values, resulting in a significantly diminished relative weight *ω*_*i*_*/*Ω. This approach allowed us to identify deviated atoms in EMD-11103 (Figure 1 of the Supplementary Material (Tu et al., 2026)).

### 4.4. Sensitivity of Global Metrics and Resolution-Dependent Sensitivity

The sensitivity analysis presented in Figure 8 underscores the limitations of similarity indices like the Q-score. Although our RHL-derived estimates provide a statistically superior fit to the local 3D density, the resulting Q-scores showed only marginal improvements over those calculated using default parameters. This relative invariance suggests that correlation-based metrics are mathematically insensitive to the atom-specific variations captured by our model.

Crucially, the RHL model exhibits resolution-dependent sensitivity. Our analysis across a multi-map portfolio demonstrates that while the 1.25 Å map reveals systematic, statistically significant signal deviations (*p* = 9.88 *×* 10^−11^), the 2.2 Å map aligns more closely with conventional uniform-width approximations. This finding demonstrates the model’s integrity as a diagnostic tool; it provides high-precision insights only when the data resolution permits, while remaining consistent with established standards when resolution limits prevent sub-atomic differentiation.

### 4.5. Conclusion and Future Perspectives

In summary, the RHL framework offers a robust, data-adaptive method for the quantitative analysis of cryo-EM maps. By implementing an automated cross-validation procedure (Algorithms 4 and 5) to dynamically select tuning parameters *α*_*r*_ and *α*_*g*_, we have removed the need for manual calibration and ensured the model is reasonably optimized for the specific noise characteristics of each map. This approach has allowed us to reveal previously obscured signal patterns, such as the Nitrogen Anomaly, which we present as an open problem for the structural biology community.

In this study, we move beyond the standard uniform-width Gaussian assumption by introducing a robust hierarchical framework for estimating atom-specific Gaussian parameters. Within this framework, signal overlap is treated as a form of data contamination, whose influence is reduced through the MDPDE weighting scheme. While the current framework employs a parsimonious isotropic Gaussian model, our results demonstrate its capacity to extract chemically meaningful features that remain inaccessible to existing pipelines. We hope this work will serve as a catalyst for the statistical community to develop more sophisticated frameworks, such as anisotropic Gaussian models that may capture even finer layers of chemical and electrostatic information in high-resolution cryo-EM data.

## Acknowledgments

The authors would like to thank the Editor, an Associate Editor, and two referees for many constructive comments and suggestions which greatly improved the quality of the paper. I-Ping Tu serves as the corresponding author for this work.

## SUPPLEMENTARY MATERIAL

### Supplement to “A Robust Hierarchical Linear Model for Cryo-EM Map Analysis”

Appendix for method section and additional numerical study.

### Analysis Source Code

The complete source code is openly available on GitHub at: https://github.com/yslianTim/rhbm_gem

## REFERENCES

Afonine, P. V., Poon, B. K., Read, R. J., Sobolev, O. V., Terwilliger, T. C., Urzhumtsev, A. and Adams, P. D. (2018). Real-space refinement in PHENIX for cryo-EM and crystallography. Acta Crystallographica Section D 74 531–544. 10.1107/S2059798318006551

Basu, A., Harris, I. R., Hjort, N. L. and Jones, M. (1998). Robust and efficient estimation by minimising a density power divergence. Biometrika 85 549–559.

Chen, T.-L., Hsieh, D.-N., Hung, H., Tu, I.-P., Wu, P.-S., Wu, Y.-M., Chang, W.-H. and Huang, S.-Y. (2014). γ-SUP: A clustering algorithm for cryo-electron microscopy images of asymmetric particles. The Annals of Applied Statistics 8 259–285.

Cheng, Y. (1995). Mean shift, mode seeking, and clustering. IEEE Transactions on Pattern Analysis and Machine Intelligence 17 790–799. 10.1109/34.400568

Cianfrocco, M. A., Lahiri, I., DiMaio, F. and Leschziner, A. E. (2018). cryoem-cloud-tools: A software platform to deploy and manage cryo-EM jobs in the cloud. Journal of Structural Biology 203 230–235. 10.1016/j.jsb.2018.05.014

Fujisawa, H. and Eguchi, S. (2008). Robust parameter estimation with a small bias against heavy contamination. J. Multivariate Anal. 99 2053–2081.

Gelman, A., Carlin, J. B., Stern, H. S. and Rubin, D. B. (2013). Bayesian Data Analysis. Chapman and Hall/CRC.

Ghosh, A. and Basu, A. (2013). Robust estimation for independent non-homogeneous observations using density power divergence with applications to linear regression. Electronic Journal of Statistics 7 2420–2456. 10.1214/13-EJS847

Ghosh, A. and Basu, A. (2016). Robust Bayes estimation using the density power divergence. Ann Inst Stat Math 68 413–437. 10.1007/s10463-014-0499-0

Jamali, K., KÄll, L., Zhang, R., Brown, A., Kimaniu, D. and Scheres, S. H. W. (2024). Automated model building and protein identification in cryo-EM maps. Nature 628 450–457. 10.1038/s41586-024-07215-4

Kirkland, E. J. (2020). Calculation of Images of Thin Specimens ; Theory of Calculation of Images of Thick Specimens In Advanced Computing in Electron Microscopy 99-195. Springer International Publishing, Cham. 10.1007/978-3-030-33260-0

KÜhlbrandt, W. (2014). The resolution revolution. Science 343 1443–1444. DOI:10.1126/science.1251652

Marques, M. A., Purdy, M. D. and Yeager, M. (2019). Cryo-EM maps are full of potential. Current Opinion in Structural Biology 58 214–223. 10.1016/j.sbi.2019.04.006

Murshudov, G. N., SkubÁk, P., Lebedev, A. A., Pannu, N. S., Steiner, R. A., Nicholls, R. A., Winn, M. D., Long, F. and Vagin, A. A. (2011). REFMAC5 for the refinement of macromolecular crystal structures. Acta Crystallographica Section D 67 355–367. 10.1107/S0907444911001314

Nirwal, S., Czarnocki-Cieciura, M., Zajko, W., Skowronek, K., Szczepanowski, R. H. and Nowotny, M. (2025). Structural snapshots of the mechanism of ATP-dependent DNA damage recognition by UvrA. Nature Communications 17. 10.1038/s41467-025-67075-y

Pintilie, G., Zhang, K., Su, Z., Li, S., Schmid, M. F. and Chiu, W. (2020). Measurement of atom resolvability in cryo-EM maps with Q-scores. Nature methods 17 328–334. 10.1038/s41592-020-0731-1

Rosenthal, P. B. and Henderson, R. (2003). Optimal determination of particle orientation, absolute hand, and contrast loss in single-particle electron cryomicroscopy. Journal of Molecular Biology 333 721–745. DOI:10.1016/j.jmb.2003.07.013

Saibil, H. (2017). Blob-ology and biology of cryo-EM:. BMC Biology 15 87. DOI:10.1186/s12915-017-0417-z

Tu, I.-P., Zheng, S.-C., Lien, Y.-H., Lin, S.-H., Lin, P.-C. and Chang, W.-H. (2026). Supplement to “A Robust Hierarchical Linear Model for Cryo-EM Map Analysis.”.

Wang, M., Jiang, L., Jian, R., Chan, J. Y., Liu, Q., Snyder, M. P. and Tang, H. (2021). RobNorm: model-based robust normalization method for labeled quantitative mass spectrometry proteomics data. Bioinformatics 37 815–821. 10.1093/bioinformatics/btaa904

Windham, M. P. (1995). Robustifying Model Fitting. Journal of the Royal Statistical Society. Series B (Methodological) 57 599–609. 10.1093/bioinformatics/btaa904

Yip, K. M., Fischer, N., Paknia, E., Chari, A. and Stark, H. (2020). Atomic-resolution protein structure determination by cryo-EM. Nature 587 157–161. 10.1038/s41586-020-2833-4

