## Supplement for "A Robust Hierarchical Linear Model for Cryo-EM Map Analysis"

### APPENDIX A: FOR METHOD SECTION

**A.1. Statistical Model.** The physics model for the cryo-EM map value on the $j^{th}$ sampling grid point of the $i^{th}$ atom is $A_i\phi(r_{ij};\tau_i^2)$ where $A_i$ is the amplitude parameter for the $i^{th}$ atom, $\phi(r_{ij};\tau_i^2)$ is the 3D Normal density function for the $i^{th}$ atom with mean 0 and width parameter $\tau_i$ and $r_{ij}$ is the distance from the $j^{th}$ sampling grid point to the $i^{th}$ atom center. After taking log transformation, we build a linear regression model as

$$
\begin{aligned}
y_{ij} =& \log[\phi_(r_{ij};\tau_i^2)A_i] + \epsilon_{ij} \\
=& \log[(2\pi\tau_i^2)^{\frac{-3}{2}} \exp\left(-\frac{1}{2\tau_i^2} r_{ij}^2\right) A_i] + \epsilon_{ij} \\
=& \left[-\frac{3}{2}\log(2\pi\tau_i^2) + \log A_i\right] + \frac{1}{\tau_i^2}\left[-\frac{1}{2}r_{ij}^2\right] + \epsilon_{ij} \\
=& \boldsymbol{x}_{ij}^\top \boldsymbol{\beta}_i + \epsilon_{ij}, \qquad (1)
\end{aligned}
$$

where $1 \leq j \leq n_i$ with $n_i$ being the sampling size for the $i^{th}$ atom, $1 \leq i \leq I$, $\boldsymbol{\beta}_i = \left(\log A_i - \frac{3}{2}\log(2\pi\tau_i^2),\ \frac{1}{\tau_i^2}\right)^\top$, $\boldsymbol{x}_{ij} = \left(1,\ -\frac{1}{2}r_{ij}^2\right)^\top$ with $r_{ij}$ being the distance of the $j^{th}$ sampling grid point to the atom center. We further let $\boldsymbol{X}_i = (\boldsymbol{x}_{i1}, \ldots, \boldsymbol{x}_{in_i})^\top$ to have the vector form for this linear model as

$$
\boldsymbol{y}_i = \boldsymbol{X}_i\boldsymbol{\beta}_i + \boldsymbol{\epsilon}_i,
$$

that the length of $\boldsymbol{y}_i$ is $n_i$.

**A.2. Map Value Sampling Scheme.** Cryo-EM maps are defined on three-dimensional regular voxel grids, so evaluating map values at arbitrary spatial locations typically requires interpolation. To obtain the map values corresponding to each atom's local environment, we apply the tricubic interpolation scheme described in Ref. Afonine et al. (2018) (Appendix B), which is commonly used for accurate evaluation of cryo-EM map values.

For each atom $i$ with position vector $\boldsymbol{r}_i$, we generate 1,500 random sampling points $\{\tilde{\boldsymbol{x}}_{ij}\}_{j=1}^{1500}$ distributed within a sphere of radius 1.5 Å centered at $\boldsymbol{r}_i$. The corresponding interpolated map values $\{\tilde{y}_{ij}\}_{j=1}^{1500}$ are obtained by evaluating the cryo-EM map at $\{\tilde{\boldsymbol{x}}_{ij}\}_{j=1}^{1500}$ using tricubic interpolation. The resulting pairs $\{(\tilde{\boldsymbol{x}}_{ij}, \tilde{y}_{ij})\}_{j=1}^{1500}$ serve as observations for estimating the model parameters of atom $i$.

*Keywords and phrases.* cryo-EM map, atomic model, robust estimate, hierarchical linear model, heteroscedasticity, minimum density power divergence estimate, PDB, EMDB.

To sample spatial locations in the vicinity of a reference point $\boldsymbol{r} \in \mathbb{R}^3$, we generate random directions uniformly over the unit sphere together with radial distances sampled within a specified range. Specifically, each sample is obtained by independently drawing

$$R \sim \mathcal{U}(0, 1.5), \qquad \phi \sim \mathcal{U}(0, 2\pi), \qquad U \sim \mathcal{U}(-1, 1),$$

where $R$ denotes the radial distance from $\boldsymbol{r}$, $\phi$ is the azimuthal angle, and $U$ corresponds to the cosine of the polar angle.

Given $U = \cos\theta$, we obtain

$$\theta = \arccos(U), \qquad \sin\theta = \sqrt{1 - U^2}.$$

A unit direction vector in spherical coordinates is then given by

$$\hat{\mathbf{n}}(\theta, \phi) = \begin{bmatrix} \sin\theta\cos\phi \\ \sin\theta\sin\phi \\ \cos\theta \end{bmatrix}.$$

The corresponding sampling location is obtained as

$$\tilde{\boldsymbol{x}} = \boldsymbol{r} + R\,\hat{\mathbf{n}}(\theta, \phi).$$

Sampling $\phi$ uniformly on $[0, 2\pi)$ and $U = \cos\theta$ uniformly on $[-1, 1]$ ensures isotropic sampling of directions on the sphere. In contrast, sampling $R$ uniformly on $[0, 1.5]$ does *not* yield a uniform spatial density within the spherical volume. Because the spherical volume element scales as $R^2\, dR$, uniform sampling of $R$ leads to a higher density of samples near the center and a lower density near the surface. This choice is intentional: interior regions are less affected by neighboring bonded atoms, and placing more samples near the center improves the reliability of local signal estimation for the target atom.

**A.3. Marginal Likelihood Estimates.** The marginal likelihood for $\boldsymbol{\mu}$ given $\boldsymbol{y}_1, \ldots, \boldsymbol{y}_I$ based on Equations (3) and (4) in Tu et al. (2026) is as follows.

$$Lik\left(\boldsymbol{\mu}|\boldsymbol{y}_1, \ldots, \boldsymbol{y}_I\right) = f\left(\boldsymbol{y}_1, \ldots, \boldsymbol{y}_I|\boldsymbol{\mu}\right)$$

$$= \int f\left(\boldsymbol{y}_1|\boldsymbol{\beta}_1\right) \cdots f\left(\boldsymbol{y}_I|\boldsymbol{\beta}_I\right) f\left(\boldsymbol{\beta}_1, \ldots, \boldsymbol{\beta}_I|\boldsymbol{\mu}\right) d\boldsymbol{\beta}_1 \cdots\, d\boldsymbol{\beta}_I$$

$$\propto \prod_{i=1}^{I} \left( \int e^{-\frac{1}{2}(\boldsymbol{y}_i - \boldsymbol{X}_i\boldsymbol{\beta}_i)^\top \boldsymbol{\Sigma}_{\boldsymbol{y}_i}^{-1}(\boldsymbol{y}_i - \boldsymbol{X}_i\boldsymbol{\beta}_i)} \cdot e^{-\frac{1}{2}(\boldsymbol{\beta}_i - \boldsymbol{\mu})^\top \boldsymbol{\Lambda}_i^{-1}(\boldsymbol{\beta}_i - \boldsymbol{\mu})} d\boldsymbol{\beta}_i \right)$$

$$\propto \prod_{i=1}^{I} \left( \int e^{-\frac{1}{2}(\boldsymbol{\beta}_i - \tilde{\boldsymbol{\beta}}_i)^\top \tilde{\boldsymbol{\Sigma}}_i^{-1}(\boldsymbol{\beta}_i - \tilde{\boldsymbol{\beta}}_i)} d\boldsymbol{\beta}_i \cdot \exp\left\{ -\frac{1}{2}\boldsymbol{y}_i^\top \boldsymbol{\Sigma}_{\boldsymbol{y}_i}^{-1}\boldsymbol{y}_i - \frac{1}{2}\boldsymbol{\mu}^\top \boldsymbol{\Lambda}_i^{-1}\boldsymbol{\mu} + \frac{1}{2}\tilde{\boldsymbol{\beta}}_i^\top \tilde{\boldsymbol{\Sigma}}_i^{-1}\tilde{\boldsymbol{\beta}}_i \right\} \right)$$

$$\propto \prod_{i=1}^{I} \left( \exp\left\{ -\frac{1}{2}\boldsymbol{\mu}^\top \boldsymbol{\Lambda}_i^{-1}\boldsymbol{\mu} + \frac{1}{2}\tilde{\boldsymbol{\beta}}_i^\top \tilde{\boldsymbol{\Sigma}}_i^{-1}\tilde{\boldsymbol{\beta}}_i \right\} \right)$$

(2)

$$\propto \prod_{i=1}^{I} \left( \exp\left\{ -\frac{1}{2}\boldsymbol{\mu}^\top \left( \boldsymbol{\Lambda}_i^{-1} - \boldsymbol{\Lambda}_i^{-1}\tilde{\boldsymbol{\Sigma}}_i\boldsymbol{\Lambda}_i^{-1} \right)\boldsymbol{\mu} + \boldsymbol{\mu}^\top \left[ \boldsymbol{\Lambda}_i^{-1}\tilde{\boldsymbol{\Sigma}}_i \left( \boldsymbol{X}_i^\top \boldsymbol{\Sigma}_{\boldsymbol{y}_i}^{-1}\boldsymbol{y}_i \right) \right] \right\} \right),$$

where $\tilde{\boldsymbol{\beta}}_i = \tilde{\boldsymbol{\Sigma}}_i \left( \boldsymbol{X}_i^\top \boldsymbol{\Sigma}_{\boldsymbol{y}_i}^{-1}\boldsymbol{y}_i + \boldsymbol{\Lambda}_i^{-1}\boldsymbol{\mu} \right)$ and $\tilde{\boldsymbol{\Sigma}}_i = \left( \boldsymbol{X}_i^\top \boldsymbol{\Sigma}_{\boldsymbol{y}_i}^{-1}\boldsymbol{X}_i + \boldsymbol{\Lambda}_i^{-1} \right)^{-1}$.

Let $Q(\boldsymbol{\mu}) = \sum_{i=1}^{I} -\frac{1}{2}\boldsymbol{\mu}^\top \left(\boldsymbol{\Lambda}_i^{-1} - \boldsymbol{\Lambda}_i^{-1}\tilde{\boldsymbol{\Sigma}}_i\boldsymbol{\Lambda}_i^{-1}\right)\boldsymbol{\mu} + \boldsymbol{\mu}^\top \left[\boldsymbol{\Lambda}_i^{-1}\tilde{\boldsymbol{\Sigma}}_i \left(\boldsymbol{X}_i^\top \boldsymbol{\Sigma}_{\boldsymbol{y}_i}^{-1}\boldsymbol{y}_i\right)\right]$, then taking the derivative of $Q(\boldsymbol{\mu})$, we have

$$\frac{dQ(\boldsymbol{\mu})}{d\boldsymbol{\mu}} = \sum_{i=1}^{I} -\tilde{\boldsymbol{\Lambda}}_i^{-1}\boldsymbol{\mu} + \boldsymbol{\Lambda}_i^{-1}\tilde{\boldsymbol{\Sigma}}_i\left(\boldsymbol{X}_i^\top \boldsymbol{\Sigma}_{\boldsymbol{y}_i}^{-1}\boldsymbol{y}_i\right),$$

where

$$\begin{aligned}
\tilde{\boldsymbol{\Lambda}}_i^{-1} &= \boldsymbol{\Lambda}_i^{-1} - \boldsymbol{\Lambda}_i^{-1}\tilde{\boldsymbol{\Sigma}}_i\boldsymbol{\Lambda}_i^{-1} \\
&= \boldsymbol{\Lambda}_i^{-1}(\boldsymbol{\Lambda}_i - \left(\boldsymbol{X}_i^\top \boldsymbol{\Sigma}_{\boldsymbol{y}_i}^{-1}\boldsymbol{X}_i + \boldsymbol{\Lambda}_i^{-1}\right)^{-1})\boldsymbol{\Lambda}_i^{-1} \\
&= \boldsymbol{\Lambda}_i^{-1}\left(\boldsymbol{X}_i^\top \boldsymbol{\Sigma}_{\boldsymbol{y}_i}^{-1}\boldsymbol{X}_i + \boldsymbol{\Lambda}_i^{-1}\right)^{-1}\left((\boldsymbol{X}_i^\top \boldsymbol{\Sigma}_{\boldsymbol{y}_i}^{-1}\boldsymbol{X}_i + \boldsymbol{\Lambda}_i^{-1})\boldsymbol{\Lambda}_i - I_2\right)\boldsymbol{\Lambda}_i^{-1} \\
&= \boldsymbol{\Lambda}_i^{-1}\left(\boldsymbol{X}_i^\top \boldsymbol{\Sigma}_{\boldsymbol{y}_i}^{-1}\boldsymbol{X}_i + \boldsymbol{\Lambda}_i^{-1}\right)^{-1}\left((\boldsymbol{X}_i^\top \boldsymbol{\Sigma}_{\boldsymbol{y}_i}^{-1}\boldsymbol{X}_i\boldsymbol{\Lambda}_i + I_2) - I_2\right)\boldsymbol{\Lambda}_i^{-1} \\
&= \boldsymbol{\Lambda}_i^{-1}\left(\boldsymbol{X}_i^\top \boldsymbol{\Sigma}_{\boldsymbol{y}_i}^{-1}\boldsymbol{X}_i + \boldsymbol{\Lambda}_i^{-1}\right)^{-1}(\boldsymbol{X}_i^\top \boldsymbol{\Sigma}_{\boldsymbol{y}_i}^{-1}\boldsymbol{X}_i) \\
&= \boldsymbol{\Lambda}_i^{-1}\tilde{\boldsymbol{\Sigma}}_i(\boldsymbol{X}_i^\top \boldsymbol{\Sigma}_{\boldsymbol{y}_i}^{-1}\boldsymbol{X}_i). \qquad (3)
\end{aligned}$$

Then, the MLE by solving $\frac{dQ(\boldsymbol{\mu})}{d\boldsymbol{\mu}} = 0$ is as follows.

$$\begin{aligned}
\hat{\boldsymbol{\mu}} &= \left[\sum_{i=1}^{I}\tilde{\boldsymbol{\Lambda}}_i^{-1}\right]^{-1}\left[\sum_{i=1}^{I}\boldsymbol{\Lambda}_i^{-1}\tilde{\boldsymbol{\Sigma}}_i\left(\boldsymbol{X}_i^\top \boldsymbol{\Sigma}_{\boldsymbol{y}_i}^{-1}\boldsymbol{y}_i\right)\right] \\
&= \left[\sum_{i=1}^{I}\boldsymbol{\Lambda}_i^{-1}\tilde{\boldsymbol{\Sigma}}_i\left(\boldsymbol{X}_i^\top \boldsymbol{\Sigma}_{\boldsymbol{y}_i}^{-1}\boldsymbol{X}_i\right)\right]^{-1}\left[\sum_{i=1}^{I}\boldsymbol{\Lambda}_i^{-1}\tilde{\boldsymbol{\Sigma}}_i\left(\boldsymbol{X}_i^\top \boldsymbol{\Sigma}_{\boldsymbol{y}_i}^{-1}\boldsymbol{y}_i\right)\right] \qquad (4)
\end{aligned}$$

**A.4. Weights.**

A.4.1. *Weights for Liner Regression.* To estimate $\sigma^2$, we have $u_\sigma(z_j) = \frac{\partial}{\partial\sigma}\log(f_\sigma(z_j)) = \frac{\partial}{\partial\sigma}(-z_j^2/(2\sigma^2) - \log(\sigma)) = z_j^2/\sigma^3 - 1/\sigma$.

$$\int u(z)f^{1+\alpha}(z)dz = \left(2\pi\sigma^2\right)^{\frac{-1-\alpha}{2}}\int\left(\frac{z^2}{\sigma^3} - \frac{1}{\sigma}\right)\exp\left[-\frac{(1+\alpha)z^2}{2\sigma^2}\right]dz = -\left(2\pi\sigma^2\right)^{\frac{-1-\alpha}{2}}\frac{\sqrt{2\pi}\alpha}{(1+\alpha)^{3/2}}$$

The estimating equation (9) in Tu et al. (2026) becomes

$$\frac{1}{n}\sum_{j=1}^{n}\left(\frac{z_j^2}{\sigma^2} - 1\right)\exp\left\{-\frac{\alpha z_j^2}{2\sigma^2}\right\} + \frac{\alpha}{(1+\alpha)^{3/2}} = 0, \qquad (5)$$

and the MDPDE of $\sigma^2$ is as follows.

$$\hat{\sigma}^2 = \left[\mathrm{tr}\left(\boldsymbol{W}\right) - \frac{n\alpha}{(1+\alpha)^{3/2}}\right]^{-1}\left(\boldsymbol{y} - \boldsymbol{X}\hat{\boldsymbol{\beta}}\right)^\top\boldsymbol{W}\left(\boldsymbol{y} - \boldsymbol{X}\hat{\boldsymbol{\beta}}\right) \qquad (6)$$

A.4.2. *Consistency for $\hat{\boldsymbol{\mu}}_{\text{MDPDE}}$.* Let $\boldsymbol{X}_I = \frac{1}{I}\sum_{i=1}^{I}\boldsymbol{\beta}_i f_{\boldsymbol{\mu}}^{\alpha}(\boldsymbol{\beta}_i)$ and $Y_I = \frac{1}{I}\sum_{j=1}^{I} f_{\boldsymbol{\mu}}^{\alpha}(\boldsymbol{\beta}_j)$, then we have

$$\lim_{I\to\infty}\hat{\boldsymbol{\mu}}_{\text{MDPDE}} = \lim_{I\to\infty}\frac{\sum_{i=1}^{I}\omega_i\boldsymbol{\beta}_i}{\sum_{i=1}^{I}\omega_i} = \lim_{I\to\infty}\frac{\frac{1}{I}\sum_{i=1}^{I}\boldsymbol{\beta}_i f_{\boldsymbol{\mu}}^{\alpha}(\boldsymbol{\beta}_i)}{\frac{1}{I}\sum_{j=1}^{I} f_{\boldsymbol{\mu}}^{\alpha}(\boldsymbol{\beta}_j)} = \lim_{I\to\infty}\frac{\boldsymbol{X}_I}{Y_I}.$$

Given that

$$\lim_{I\to\infty} \boldsymbol{X}_I = E_{f_\mu}[\boldsymbol{\beta} f_{\boldsymbol{\mu}}^{\alpha}(\boldsymbol{\beta})] = \int \boldsymbol{\beta} f_{\boldsymbol{\mu}}^{1+\alpha}(\boldsymbol{\beta}) d\boldsymbol{\beta},$$

$$\lim_{I\to\infty} Y_I = E_{f_\mu}[f_{\boldsymbol{\mu}}^{\alpha}(\boldsymbol{\beta})] = \int f_{\boldsymbol{\mu}}^{1+\alpha}(\boldsymbol{\beta}) d\boldsymbol{\beta},$$

we can then apply Slutsky's Theorem which is provided in the end of this section, to get

$$\lim_{I\to\infty} \hat{\boldsymbol{\mu}}_{\text{MDPDE}} = \lim_{I\to\infty} \frac{\boldsymbol{X}_I}{Y_I} = \frac{\lim_{I\to\infty} \boldsymbol{X}_I}{\lim_{I\to\infty} Y_I} = \frac{\int \boldsymbol{\beta} f_{\boldsymbol{\mu}}^{1+\alpha}(\boldsymbol{\beta}) d\boldsymbol{\beta}}{\int f_{\boldsymbol{\mu}}^{1+\alpha}(\boldsymbol{\beta}) d\boldsymbol{\beta}} = \boldsymbol{\mu}. \tag{7}$$

**Slutsky's Theorem**

Let $X_n$ and $Y_n$ be sequences of scalar or vector or matrix random elements. If $X_n$ converges in distribution to a random element $X$ and $Y_n$ converges in probability to a constant $c$, then

$$X_n + Y_n \to X + c,$$

$$X_n * Y_n \to X * c,$$

$$X_n / Y_n \to X/c \text{ if c is invertible.}$$

A.4.3. *The MDPDE for Estimating the $p$-dimensional Covariance Matrix $\boldsymbol{\Lambda}$.* To derive the robust estimator for the group-level covariance matrix $\boldsymbol{\Lambda}$ of dimension $p \times p$, we define the score function as follows. For $\boldsymbol{\beta}_i \sim \mathcal{N}(\boldsymbol{\mu}, \boldsymbol{\Lambda})$, where $1 \le i \le I$, the estimation of $\boldsymbol{\Lambda}$ follows the MDPDE framework established in Equation (9) in Tu et al. (2026). We first derive the score function $u_{\boldsymbol{\Lambda}}(\boldsymbol{\beta})$ with respect to the elements of the covariance matrix.

The maximum likelihood score function for a specific element $\Lambda_k$ of the covariance matrix $\boldsymbol{\Lambda}$ is given by:

$$u_{\Lambda_k}(\boldsymbol{\beta}) = \frac{\partial \log f_{\boldsymbol{\Lambda}}(\boldsymbol{\beta})}{\partial \Lambda_k} = -\frac{1}{2}\frac{\partial}{\partial \Lambda_k}\left[\log|\boldsymbol{\Lambda}| + (\boldsymbol{\beta}-\boldsymbol{\mu})^\top \boldsymbol{\Lambda}^{-1}(\boldsymbol{\beta}-\boldsymbol{\mu})\right]. \tag{8}$$

We employ the matrix differentiation identities from Harville Harville (1997):

$$\begin{cases} \frac{\partial \log|\boldsymbol{F}|}{\partial x_j} = \text{tr}\left(\boldsymbol{F}^{-1}\frac{\partial \boldsymbol{F}}{\partial x_j}\right) \\ \frac{\partial \boldsymbol{F}^{-1}}{\partial x_j} = -\boldsymbol{F}^{-1}\frac{\partial \boldsymbol{F}}{\partial x_j}\boldsymbol{F}^{-1}. \end{cases} \tag{9}$$

Applying these identities to the log-likelihood of the Normal distribution, the score function with respect to the element $\Lambda_k$ is:

$$\begin{aligned} u_{\Lambda_k}(\boldsymbol{\beta}) &= -\frac{1}{2}\text{tr}\left(\boldsymbol{\Lambda}^{-1}\frac{\partial \boldsymbol{\Lambda}}{\partial \Lambda_k}\right) + \frac{1}{2}(\boldsymbol{\beta}-\boldsymbol{\mu})^\top \boldsymbol{\Lambda}^{-1}\frac{\partial \boldsymbol{\Lambda}}{\partial \Lambda_k}\boldsymbol{\Lambda}^{-1}(\boldsymbol{\beta}-\boldsymbol{\mu}) \\ &= \frac{1}{2}\text{tr}\left\{\boldsymbol{\Lambda}^{-1}(\boldsymbol{\beta}-\boldsymbol{\mu})(\boldsymbol{\beta}-\boldsymbol{\mu})^\top\boldsymbol{\Lambda}^{-1}\frac{\partial \boldsymbol{\Lambda}}{\partial \Lambda_k} - \boldsymbol{\Lambda}^{-1}\frac{\partial \boldsymbol{\Lambda}}{\partial \Lambda_k}\right\} \\ &= \frac{1}{2}\text{tr}\left\{\left[\boldsymbol{\Lambda}^{-1}(\boldsymbol{\beta}-\boldsymbol{\mu})(\boldsymbol{\beta}-\boldsymbol{\mu})^\top - \mathbf{I}_p\right]\boldsymbol{\Lambda}^{-1}\frac{\partial \boldsymbol{\Lambda}}{\partial \Lambda_k}\right\}. \end{aligned} \tag{10}$$

The resulting MDPDE estimating equation for each element $\Lambda_k$ is:

$$U_I(\Lambda_k) := \frac{1}{I}\sum_{i=1}^{I} u_{\Lambda_k}(\boldsymbol{\beta}_i) f^{\alpha_g}(\boldsymbol{\beta}_i) - \int u_{\Lambda_k}(\boldsymbol{\beta}) f^{1+\alpha_g}(\boldsymbol{\beta})\text{d}\boldsymbol{\beta} = 0, \quad \forall k = 1, \ldots, p^2. \tag{11}$$

Note that while the covariance matrix $\boldsymbol{\Lambda}$ contains only $p(p+1)/2$ unique elements due to its symmetry, we employ the $p^2$ redundant notation to simplify the indexing and the application of matrix differentiation identities.

Since the parameters $\boldsymbol{\Lambda}$ and $\Lambda_k$ are independent of $\boldsymbol{\beta}$, the integral term in Equation (11) involves the following two key integrals, evaluated over the $p$-dimensional domain:

$$\int(\boldsymbol{\beta}-\boldsymbol{\mu})(\boldsymbol{\beta}-\boldsymbol{\mu})^\top f^{1+\alpha_g}(\boldsymbol{\beta})\mathrm{d}\boldsymbol{\beta}=[(2\pi)^p|\boldsymbol{\Lambda}|]^{-\alpha_g/2}(1+\alpha_g)^{-1-\frac{p}{2}}\boldsymbol{\Lambda}, \tag{12}$$

$$\int \mathbf{I}_p f^{1+\alpha_g}(\boldsymbol{\beta})\mathrm{d}\boldsymbol{\beta}=[(2\pi)^p|\boldsymbol{\Lambda}|]^{-\alpha_g/2}(1+\alpha_g)^{-p/2}\mathbf{I}_p. \tag{13}$$

Substituting Equations (12) and (13) into the integral component of the estimating equation (11), we obtain:

$$\begin{aligned}
&\int u_{\Lambda_k}(\boldsymbol{\beta})f^{1+\alpha_g}(\boldsymbol{\beta})\mathrm{d}\boldsymbol{\beta}\\
&=\frac{1}{2}\mathrm{tr}\left\{\left[\boldsymbol{\Lambda}^{-1}\int(\boldsymbol{\beta}-\boldsymbol{\mu})(\boldsymbol{\beta}-\boldsymbol{\mu})^\top f^{1+\alpha_g}(\boldsymbol{\beta})\mathrm{d}\boldsymbol{\beta}-\int \mathbf{I}_p f^{1+\alpha_g}(\boldsymbol{\beta})\mathrm{d}\boldsymbol{\beta}\right]\boldsymbol{\Lambda}^{-1}\frac{\partial\boldsymbol{\Lambda}}{\partial\Lambda_k}\right\}\\
&=\frac{1}{2}[(2\pi)^p|\boldsymbol{\Lambda}|]^{-\alpha_g/2}\,\mathrm{tr}\left\{\left[(1+\alpha_g)^{-1-\frac{p}{2}}\mathbf{I}_p-(1+\alpha_g)^{-p/2}\mathbf{I}_p\right]\boldsymbol{\Lambda}^{-1}\frac{\partial\boldsymbol{\Lambda}}{\partial\Lambda_k}\right\}\\
&=-\frac{\alpha_g}{2}[(2\pi)^p|\boldsymbol{\Lambda}|]^{-\alpha_g/2}(1+\alpha_g)^{-1-\frac{p}{2}}\mathrm{tr}\left(\boldsymbol{\Lambda}^{-1}\frac{\partial\boldsymbol{\Lambda}}{\partial\Lambda_k}\right).
\end{aligned} \tag{14}$$

Combining the score function from Equation (10) and the integral term from Equation (14), the estimating equation for $\Lambda_k$ becomes:

$$\begin{aligned}
&\frac{1}{I}\sum_{i=1}^{I}u_{\Lambda_k}(\boldsymbol{\beta}_i)f^{\alpha_g}(\boldsymbol{\beta}_i)-\int u_{\Lambda_k}(\boldsymbol{\beta})f^{1+\alpha_g}(\boldsymbol{\beta})\mathrm{d}\boldsymbol{\beta}=0\\
&\Rightarrow\mathrm{tr}\left\{\left[\sum_{i=1}^{I}(\boldsymbol{\beta}_i-\boldsymbol{\mu})(\boldsymbol{\beta}_i-\boldsymbol{\mu})^\top\omega_i-\left(\Omega-I\alpha_g(1+\alpha_g)^{-1-\frac{p}{2}}\right)\boldsymbol{\Lambda}\right]\boldsymbol{\Lambda}^{-1}\frac{\partial\boldsymbol{\Lambda}}{\partial\Lambda_k}\boldsymbol{\Lambda}^{-1}\right\}=0,
\end{aligned} \tag{15}$$

where $\omega_i=\exp\left[-\frac{\alpha_g}{2}(\boldsymbol{\beta}_i-\boldsymbol{\mu})^\top\boldsymbol{\Lambda}^{-1}(\boldsymbol{\beta}_i-\boldsymbol{\mu})\right]$ and $\Omega=\sum_{i=1}^{I}\omega_i$. Since this equality must hold for all basis matrices $\frac{\partial\boldsymbol{\Lambda}}{\partial\Lambda_k}$, the term inside the brackets must vanish, yielding the explicit MDPDE for the unstructured covariance matrix:

$$\hat{\boldsymbol{\Lambda}}_{\mathrm{MDPDE}}=\frac{\sum_{i=1}^{I}\omega_i(\boldsymbol{\beta}_i-\boldsymbol{\mu})(\boldsymbol{\beta}_i-\boldsymbol{\mu})^\top}{\Omega-I\alpha_g(1+\alpha_g)^{-1-\frac{p}{2}}}. \tag{16}$$

This derivation confirms that the consistency factor depends directly on the dimension $p$. Unlike existing literature that often assumes a diagonal structure, Equation (3) in Tu et al. (2026) accommodates a fully unstructured $\boldsymbol{\Lambda}$, providing a robust framework for capturing correlations between all $p$ parameters in the hierarchical model.

**A.5. Influence Functions for MLE and MDPDE.** To provide a theoretical foundation for the robustness of the RHL framework, we examine the **influence function (IF)**, a key tool in robust statistics that measures the sensitivity of an estimator to an infinitesimal amount of contamination at a specific point $\tilde{\boldsymbol{x}}$. For a functional $T$ and a distribution $F$, the influence function is defined as:

$$IF(\tilde{\boldsymbol{x}};T,F) = \lim_{\epsilon\to 0} \frac{T((1-\epsilon)F + \epsilon\delta_{\tilde{\boldsymbol{x}}}) - T(F)}{\epsilon}, \tag{17}$$

where $\delta_{\tilde{\boldsymbol{x}}}$ represents a point mass at the observation $\tilde{\boldsymbol{x}}$.

A.5.1. *Influence Function of the Maximum Likelihood Estimator (MLE).* For the population mean $\boldsymbol{\mu}$ under the assumption of a Normal distribution $\mathcal{N}(\boldsymbol{\mu}, \boldsymbol{\Lambda})$, the MLE is derived from the score function $u_{\boldsymbol{\mu}}(\tilde{\boldsymbol{x}}) = \boldsymbol{\Lambda}^{-1}(\tilde{\boldsymbol{x}} - \boldsymbol{\mu})$. The influence function for the MLE is proportional to this score:

$$IF(\tilde{\boldsymbol{x}}; \text{MLE}, \Phi_{\boldsymbol{\mu}}) = \tilde{\boldsymbol{x}} - \boldsymbol{\mu}. \tag{18}$$

The linearity of this function implies that the influence of an observation increases without bound as $\tilde{\boldsymbol{x}}$ moves further from the population mean $\boldsymbol{\mu}$. Consequently, a single extreme outlier in the cryo-EM data can arbitrarily bias the resulting estimate.

A.5.2. *Influence Function of the MDPDE.* In contrast, the Minimum Density Power Divergence Estimator (MDPDE) utilizes a power parameter $\alpha > 0$ to weight the score function by the model density $f^{\alpha}_{\boldsymbol{\mu}}(\tilde{\boldsymbol{x}})$. For the mean $\boldsymbol{\mu}$ in the RHL model, the influence function is given by:

$$IF(\tilde{\boldsymbol{x}}; \text{MDPDE}, \Phi_{\boldsymbol{\mu}}) = M_{\alpha}^{-1} \left[ (\tilde{\boldsymbol{x}} - \boldsymbol{\mu}) \exp\left( -\frac{\alpha}{2} (\tilde{\boldsymbol{x}} - \boldsymbol{\mu})^{\top} \boldsymbol{\Lambda}^{-1} (\tilde{\boldsymbol{x}} - \boldsymbol{\mu}) \right) \right], \tag{19}$$

where $M_{\alpha}$ is a scalar bounded away from zero.

- **Redescending Property**: Unlike the MLE, this influence function is redescending. As the distance between the observation $\tilde{\boldsymbol{x}}$ and the mean $\boldsymbol{\mu}$ increases, the exponential term dominates, causing the influence to decay toward zero.
- **Bounded Influence**: The influence is strictly bounded, ensuring that observations far from the expected Gaussian profile (data contamination) have a negligible impact on the group-level estimates.

A.5.3. *Implications for Diagnostic Utility: The Robustness Paradox.* The redescending nature of the MDPDE influence function leads to a **"robustness paradox"**: because the estimator so effectively neutralizes the influence of extreme contamination, the influence function itself becomes a limited diagnostic for identifying those points. Since $IF(\tilde{\boldsymbol{x}}; \text{MDPDE}) \to 0$ for heavily down-weighted atoms, these points no longer contribute to the variance or bias of the final estimate. Therefore, we utilize **weight-based diagnostics**—specifically the relative weight profile $w_i/\Omega < \delta/I$—as the primary tool for identifying structural model misspecification in cryo-EM data.

As a practical application, we employ $\delta = 0.05$ to analyze the backbone atoms of EMD-11103 and PDB-6Z6U, labeling atoms that deviate significantly from the population consensus. Figure 1 illustrates that the deviant atoms (labeled in red) are predominantly distributed within flexible or solvent-exposed regions of the protein structure.

### APPENDIX B: FOR NUMERICAL STUDY

**B.1. Comparison of Robust Estimates under Different Power Parameters.** We investigate the impact of the power parameters on estimation performance. Figure 2 compares robust regression estimates with $\alpha_r = 0.1$ and $\alpha_r = 0.4$, while Figure 3 compares robust group-level estimates with $\alpha_g = 0.2$ and $\alpha_g = 0.5$, fixing $\alpha_r = 0.1$.

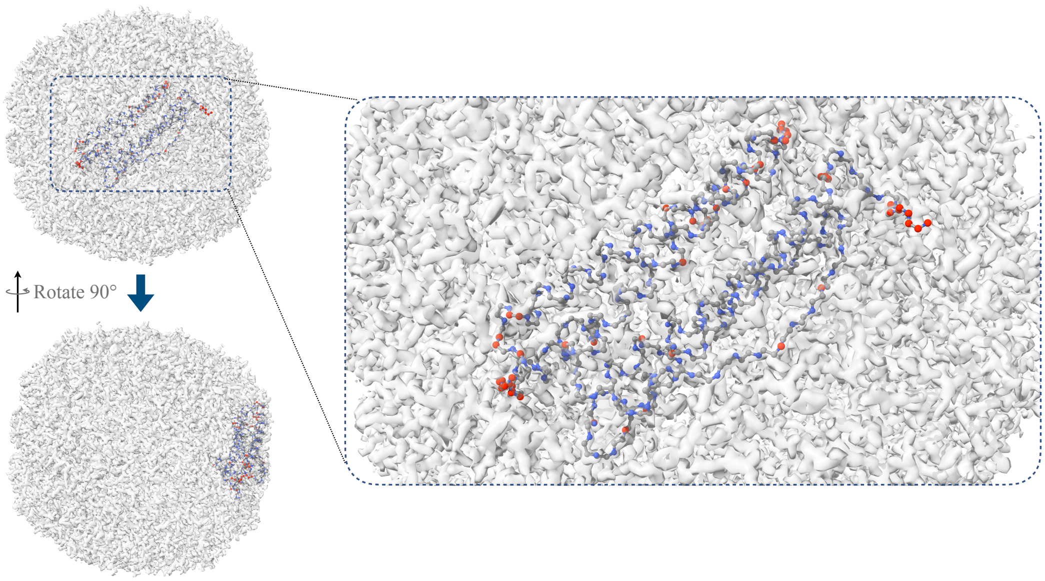


FIG 1. **Visualization of detected deviated atoms in a single unit of apoferritin of EMD-11103 and PDB-6Z6U.** *The figure shows the spatial locations of atoms identified as deviated among the backbone atoms, based on paired data from EMD-11103 (EMDB) and PDB-6Z6U. Deviated atoms are defined as atoms with weights smaller than the threshold of* $0.05/I$ *in the estimation of the group-level mean, where* $I$ *is the number of atoms in the group. For reference, an equal contribution corresponds to a weight of* $1/I$*; thus, the threshold* $0.05/I$ *highlights atoms whose contributions are substantially smaller than expected due to strong deviation from the dominant signal pattern. The transparent gray volume represents the cryo-EM map density (EMD-11103). Spheres indicate atomic positions from the corresponding model: gray for carbon atoms, blue for nitrogen atoms, and red for atoms identified as "deviated".*

Figure 2 shows that, under the highest error intensity ($\sigma_\epsilon = 0.3D_{\max}$), $\alpha_r = 0.4$ controls the bias within $10\%$ for outlier ratios up to $25\%$ for the amplitude parameter and $30\%$ for the width parameter. Increasing $\alpha_r$ improves robustness against outliers but leads to increased variability in the estimates.

Figure 3 exhibits a similar pattern. With $\alpha_g = 0.5$, the bias remains within $10\%$ for outlier ratios up to $30\%$ for the amplitude parameter and $35\%$ for the width parameter. Larger $\alpha_g$ improves robustness while increasing variability.

These results provide additional evidence for the trade-off between robustness and efficiency governed by the choice of the power parameters.

**B.2. Real Data Analysis on the paired data EMD-11103 (EMDB) and PDB-6Z6U.** Figure 4 presents one-dimensional projections of the cryo-EM map values for $C_\alpha$ across 20 amino acid types, based on paired data from EMD-11103 (EMDB) and PDB-6Z6U. Each panel corresponds to an amino acid type, labeled at the top with three bold letters. The solid curves represent averaged data values at varying distances from the atom center, indexed on the X-axis. The red dashed lines indicate robust estimates with dynamically selected tuning parameters yielding $\alpha_r = 0.1$ and $\alpha_g = 0.6$, while the blue dotted lines represent non-robust estimates with $\alpha_r = \alpha_g = 0$.

The figure clearly illustrates that when no significant deviant curve presents, the robust (red dashed) and non-robust (blue dotted) estimates align closely. However, in the presence of significant deviant curves, the robust estimate curves (red dashed line) effectively mitigates

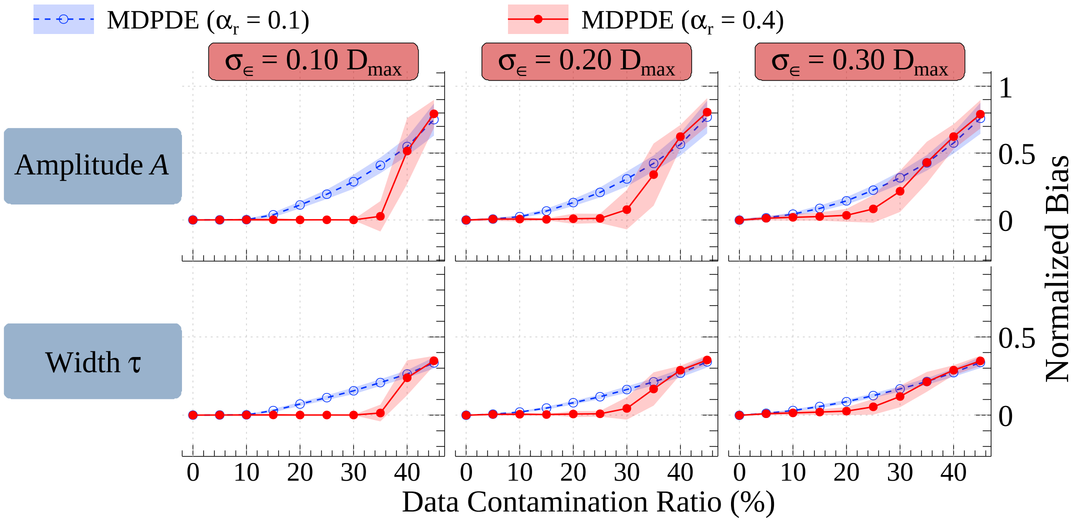


FIG 2. **Comparison on the robust regression estimates with power parameters** $\alpha_r = 0.1$ **and** $\alpha_r = 0.4$**.** *Using the simulation data illustrated in Figure 2 (Tu et al. (2026)) we evaluate estimation performance for robust regression with power parameters* $\alpha_r = 0.1$ *and* $\alpha_r = 0.4$*. Normalized bias for the amplitude parameter (top) and width parameter (bottom) are plotted against the contamination proportion. Markers denote the mean normalized bias, and colored vertical bands represent the corresponding standard deviations on both sides. The results show that* $\alpha_r = 0.4$ *provides smaller bias than* $\alpha_r = 0.1$ *across contamination levels, indicating improved robustness to outliers. However, increasing* $\alpha_r$ *also leads to greater variability in the estimates, reflecting a loss of efficiency. In the figure, estimates with* $\alpha_r = 0.1$ *are shown in solid circle and those with* $\alpha_r = 0.4$ *in open circle. The results for three error intensities in generating simulation data are represented in three columns.*

the impact from the deviant curves, while the non-robust estimate curves are obviously pulled down by the deviant curves.

**B.3. Data and Code Availability.** The data and computational tools supporting the findings of this study are available as follows:

*Data Availability.* The experimental cryo-EM maps and their corresponding atomic models analyzed in this study are publicly available in the Electron Microscopy Data Bank (EMDB) and the Protein Data Bank (PDB) under the following accession codes:

- Human Apoferritin (atomic resolution 1.25 Å)
  - PDB ID 6Z6U: https://files.rcsb.org/download/6Z6U.cif
  - EMDB ID EMD-11103: https://ftp.ebi.ac.uk/pub/databases/emdb/structures/EMD-11103/other/emd_11103_additional.map.gz
- Beta-galactosidase (near atomic resolution 2.20 Å)
  - PDB ID 6DRV: https://files.rcsb.org/download/6DRV.cif
  - EMDB ID EMD-8908: https://ftp.ebi.ac.uk/pub/databases/emdb/structures/EMD-8908/other/emd_8908_additional.map.gz
- UvrA-DNA complex (near atomic resolution 3.23 Å)
  - PDB ID 9GXM: https://files.rcsb.org/download/9GXM.cif
  - EMDB ID EMD-51667: https://ftp.ebi.ac.uk/pub/databases/emdb/structures/EMD-51667/other/emd_51667_additional_1.map.gz

*Code Availability.* All computational procedures, including the Robust Hierarchical Linear (RHL) model and the automated tuning parameter selection strategies (Algorithms 1-5

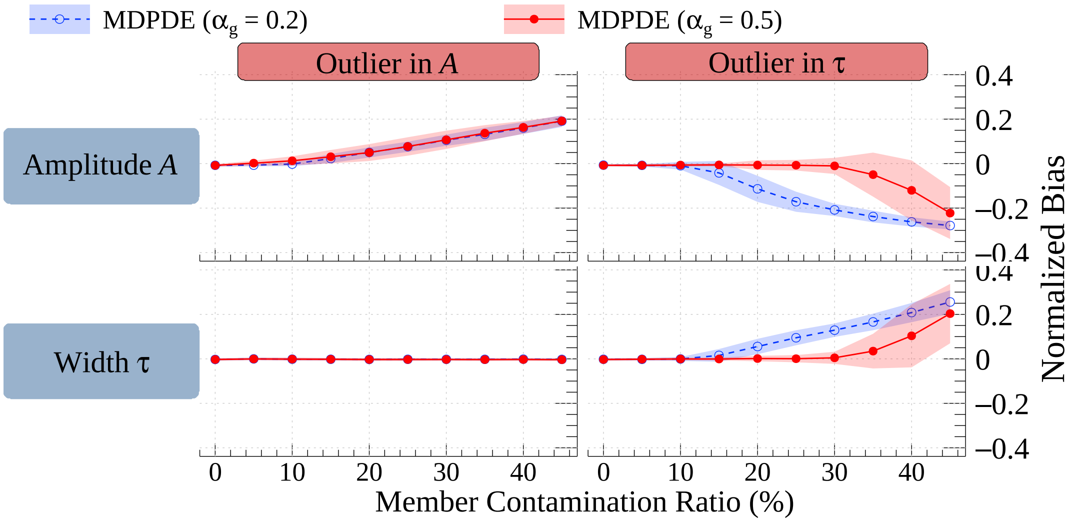


FIG 3. **Comparison of robust group-level estimates under different power parameters.** *This figure compares the normalized bias of robust group-level estimates with power parameters $\alpha_g = 0.2$ and $\alpha_g = 0.5$. Estimates with $\alpha_g = 0.2$ are shown in open circle and those with $\alpha_g = 0.5$ in solid circle. The left column corresponds to contamination introduced in the amplitude parameter, and the right column corresponds to contamination introduced in the width parameter. The coincidence of the two markers in the lower-left panel indicates that contamination in amplitude has negligible impact on width estimation for both choices of the tuning parameter in this setting. The results show that $\alpha_g = 0.5$ generally yields smaller bias than $\alpha_g = 0.2$, or performs comparably across contamination levels, indicating improved robustness to outliers. Similar to Figure 2, increasing $\alpha_g$ enhances robustness but also introduces greater variability in the estimates, reflecting a loss of efficiency.*

in Tu et al. (2026)), are implemented in C++. The complete source code, including preprocessing scripts for tricubic interpolation and simulation routines used for the performance benchmarks, is openly available on GitHub at: https://github.com/yslianTim/rhbm_gem.

### REFERENCES

AFONINE, P. V., POON, B. K., READ, R. J., SOBOLEV, O. V., TERWILLIGER, T. C., URZHUMTSEV, A. and ADAMS, P. D. (2018). Real-space refinement in PHENIX for cryo-EM and crystallography. *Acta Crystallographica Section D* **74** 531–544. https://doi.org/10.1107/S2059798318006551

HARVILLE, D. A. (1997). *Matrix Differentiation* In *Matrix Algebra From a Statistician's Perspective* 289–335. Springer New York, New York, NY. https://doi.org/10.1007/0-387-22677-X_15

TU, I.-P., ZHENG, S.-C., LIEN, Y.-H., LIN, S.-H., LIN, P.-C. and CHANG, W.-H. (2026). A Robust Hierarchical Linear Model for Cryo-EM Map Analysis. *Annals of Applied Statistics*.

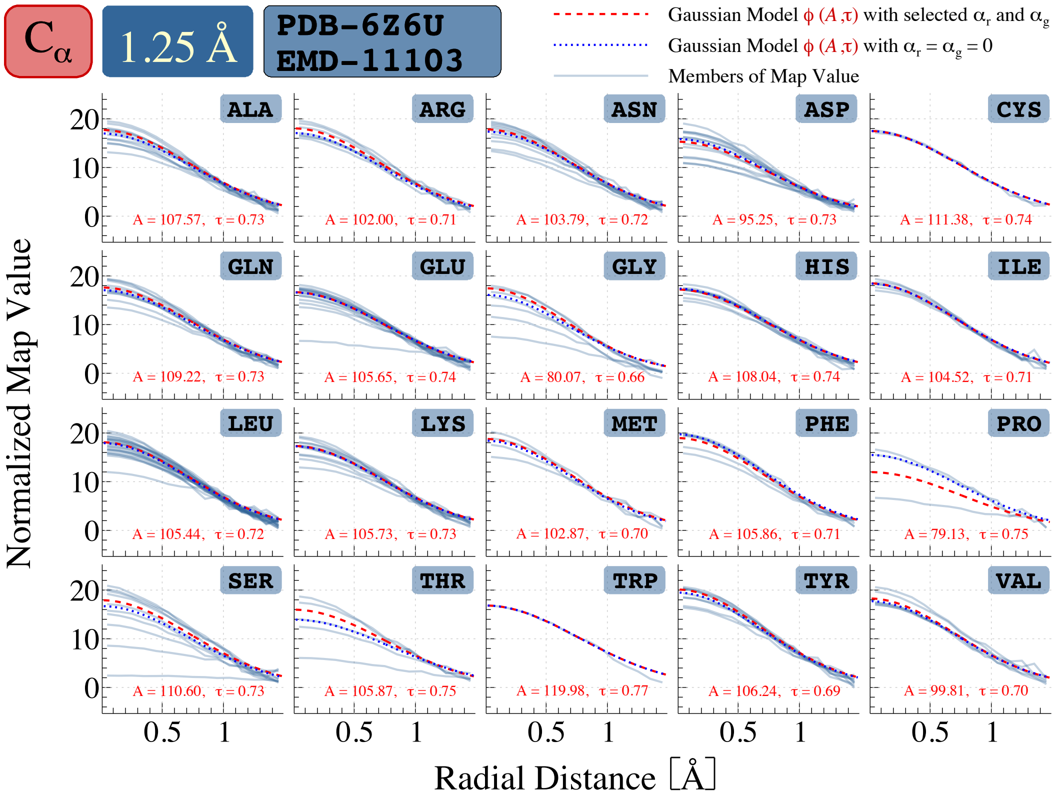


FIG 4. **Summary of real data analysis.** *The figure presents one-dimensional radial profiles of cryo-EM map values for $C_\alpha$ atoms across the 20 amino acid types, using paired data from EMD-11103 (EMDB) and PDB-6Z6U. Each panel corresponds to one amino acid type, labeled by its three-letter code. Solid curves represent the empirical data, obtained by taking the median of map values across grid points at equal distances from the atomic center, with distance shown on the x-axis. The red dashed curves correspond to the fitted signals from the robust estimator with tuning parameters selected by Algorithms 4 and 5 in Tu et al. (2026), while the blue dotted curves represent the non-robust estimator with $\alpha_r = \alpha_g = 0$.*
